# Bleaching resistant corals retain heat tolerance following acclimatization to environmentally distinct reefs

**DOI:** 10.1101/2020.09.25.314203

**Authors:** Katie L. Barott, Ariana S. Huffmyer, Jennifer M. Davidson, Elizabeth A. Lenz, Shayle B. Matsuda, Joshua R. Hancock, Teegan Innis, Crawford Drury, Hollie M. Putnam, Ruth D. Gates

**Affiliations:** Department of Biology, University of Pennsylvania, Philadelphia, PA 19104 USA; Hawai‘i Institute of Marine Biology, University of Hawai‘i, Mānoa, HI 96744 USA; Department of Biological Sciences, University of Rhode Island, Kingston, RI 02881 USA; Sea Grant College Program, University of Hawai‘i at Mānoa, HI, 96822 USA

**Keywords:** climate change, coral bleaching, coral-algal symbiosis, assisted gene flow, assisted evolution, ocean acidification, ocean warming

## Abstract

Urgent action is needed to prevent the demise of coral reefs as the climate crisis leads to an increasingly warmer and more acidic ocean. Propagating climate change resistant corals to restore degraded reefs is one promising strategy; however, empirical evidence is needed to determine if resistance is retained following transplantation within or beyond a coral’s natal reef. Here we assessed the performance of bleaching-resistant individuals of two coral species following reciprocal transplantation between environmentally distinct reefs (low *vs* high diel variability) to determine if stress resistance is retained following transplantation. Critically, transplantation to either environment had no influence on coral bleaching resistance, indicating that this trait was relatively fixed and is thus a useful metric for selecting corals for reef restoration within their native range. In contrast, growth was highly plastic, and native performance was not predictive of performance in the novel environment. Coral metabolism was also plastic, with cross transplants of both species matching the performance of native corals at both reefs within three months. Coral physiology (autotrophy, heterotrophy, and metabolism) and overall fitness (survival, growth, and reproduction) were higher at the reef with higher flow and fluctuations in diel pH and dissolved oxygen, and did not differ between native corals and cross-transplants. Conversely, cross-transplants at the low-variability reef had higher fitness than native corals, thus increasing overall fitness of the recipient population. This experiment was conducted during a non-bleaching year, which suggests that introduction of these bleaching-resistant individuals will provide even greater fitness benefits to recipient populations during bleaching years. In summary, this study demonstrates that propagating and transplanting bleaching-resistant corals can elevate the resistance of coral populations to ocean warming while simultaneously maintaining reef function as the climate crisis worsens.

## Introduction

The global climate crisis is threatening the survival of coral reef ecosystems around the world. As climate change increases the temperature of the world’s oceans (Johnson and Lyman 2020), marine heatwaves are becoming increasingly frequent (Frölicher et al. 2018) and leading to widespread coral bleaching (Hughes, Anderson, et al. 2018), a heat stress response where the coral-algal symbiosis breaks down and the algae (dinoflagellates in the family Symbiodiniaceae) are expelled from the host (Jokiel 2004; Oakley and Davy 2018). This dysbiosis has a myriad of negative consequences, ranging from declines in coral growth and reproduction to extensive coral mortality (Ward et al. 2000; Baird and Marshall 2002; Baker et al. 2008; Hughes, Kerry, Baird, et al. 2019; Hughes, Kerry, et al. 2018). These bleaching-associated outcomes affect the function of the entire reef ecosystem, as coral biomineralization is necessary to build and maintain the physical framework that is required to support the immense biodiversity typical of a healthy coral reef (Fordyce et al. 2019; Leggat et al. 2019; Hughes, Kerry, Connolly, et al. 2019). Deterioration of the reef structure is also being exacerbated by the other climate change stressor, ocean acidification (Doney et al. 2009), which has also led to declines in net ecosystem calcification (Eyre et al. 2018; Andersson and Gledhill 2013; Albright et al. 2016). An important ongoing question is whether coral populations have the capacity to acclimatize or adapt to these two climate change stressors fast enough to avoid catastrophic losses (Edmunds and Gates 2008), and whether human intervention can enhance this process to help corals keep pace with a rapidly changing environment (Van Oppen et al. 2015). Encouragingly, there is evidence that coral populations are becoming more resistant to bleaching during heat stress (Sully et al. 2019; Coles et al. 2018). However, this nominal improvement may be coming at the expense of certain species, as only the more tolerant taxa remain following the selective sieve of major bleaching mortality events (Hughes, Kerry, et al. 2018; Loya et al. 2001; McClanahan 2004; Edmunds 2018).

Action is clearly needed to mitigate widespread mortality of coral reefs predicted over the next century given business as usual carbon emissions. There has been a surge in discussions over the last few years on the implementation of adaptive management strategies such as selective propagation of climate change resistant corals (e.g. via assisted gene flow, selective breeding) to prevent the extinction of reefs and species (Van Oppen et al. 2015, 2017; Anthony et al. 2015; National Academies of Sciences et al. 2019; Anthony et al. 2020). Propagation of individuals with desired phenotypes (e.g. rapid growth, bleaching resistance) for coral reef recovery and restoration is a promising approach; however, the utility of these scientifically informed efforts depends not only on the survival of coral transplants, but also the retention of selected traits (e.g. rapid growth, bleaching resistance) following transplantation to novel physicochemical and ecological conditions and their integration into the population. Determining the feasibility of these approaches therefore requires improved knowledge of the fundamental mechanisms of coral acclimatization, since we do not know if, or for how long, these phenotypes are retained following exposure to novel environmental regimes within or across generations. Rigorous experimental evaluation is therefore needed to address this question, and the results of which are important not only for restoration, but also for understanding the capacity for coral populations to withstand rapid environmental change resulting from anthropogenic activities.

A first step in testing whether bleaching resistance is the result of local adaptation or acclimatization is to identify bleaching resistant individuals with higher temperature thresholds for bleaching within a population. These corals are often found in locations with higher mean temperatures (e.g. shallow inshore reefs with restricted water flow, (Jokiel and Brown 2004; Woesik et al. 2012; Castillo and Helmuth 2005), or those with larger magnitude or higher frequency fluctuations in temperature than surrounding reefs (Oliver and Palumbi 2011; Palumbi et al. 2014; Schoepf et al. 2015; Safaie et al. 2018), though not always (Klepac and Barshis 2020). Reefs with conditions that promote these local threshold maxima are likely excellent resources for selecting the most bleaching resistant genets of the various species found in a region, but only if elevated heat tolerance is retained when environmental conditions change. It is therefore critical to understand the biological mechanisms by which these environmental conditions influence coral bleaching thresholds, either by acclimatization or adaptation, as these mechanisms influence the persistence of adaptive traits through time and space (Drury 2020). For example, seasonal acclimatization can temporarily elevate local bleaching thresholds when peak temperatures are preceded by brief sub-bleaching heat pulses, essentially priming corals to tolerate subsequent heat stress (Ainsworth et al. 2016; Sully et al. 2019), though it appears such intra and cross-generational priming benefits can be temporary (Putnam et al. 2020). Coral bleaching events provide an opportunity to identify bleaching resistant individuals within a population, and have the advantage of allowing assessment of relative performance between individuals of the same species, in a natural context.

Here, we identified bleaching resistant individuals (i.e. corals that did not bleach during the second of two coral bleaching events that occurred in the span of two years in the Main Hawaiian Islands in 2015; Matsuda et al. 2020) of two important reef-building species, *Montipora capitata* and *Porites compressa*. After monitoring these corals for one year following the bleaching event, we tested the effects of acclimatization to a novel physicochemical environment on their bleaching resistance and fitness by reciprocally transplanting ramets of each colony between two patch reefs in Kāne‘ohe Bay, O‘ahu, Hawai‘i with contrasting environmental conditions. In addition, the physiological plasticity of each species was examined by measuring coral survival, growth, metabolism, tissue energetics, and feeding rates in their native vs. cross-transplanted environments at 3- and 6-months post-transplantation. These experiments are a critical step towards understanding the biological basis and utility of selecting and propagating climate change resistant corals for enhancing coral reef resilience to climate change.

## Materials and Methods

### Experimental Design

#### Site selection and characterization

Kāne‘ohe Bay contains a network of coral-dominated fringing and patch reefs (Fig. 1A). These reefs are protected from wave action by a barrier reef that generates a gradient of seawater residence times (Lowe et al. 2009) and thus a spatial gradient in physicochemical conditions (e.g. magnitude of diel pH and pCO2 fluctuations; (Massaro et al. 2012; Drupp et al. 2013; Page et al. 2018). Here we targeted two patch reefs representing distinct physicochemical conditions: 1) an Inner Lagoon (IL) reef (21.4343°N, 157.7991°W) with a relatively low-flow and stable pH environment, and 2) an Outer Lagoon (OL) reef (21.4516°N, 157.7966°W) with a relatively high-flow and variable pH environment (Fig. 1A). In addition to differences in pH and flow, the IL reef is located nearshore (0.75 km) and is thus exposed to greater terrestrial influence than the OL reef (1.6 km from shore; Fig. 1A).

**Figure 1.**
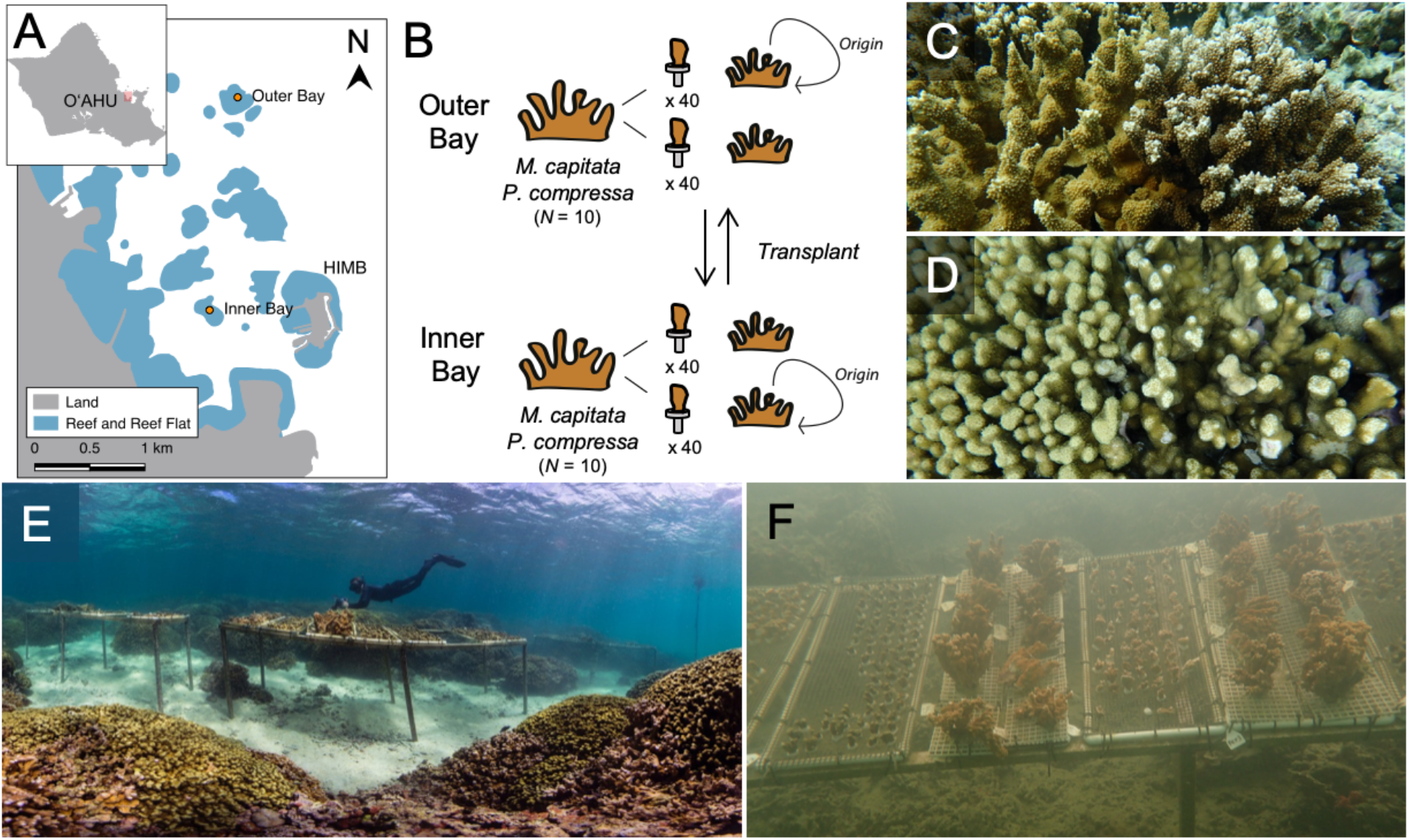
Overview of the experimental setup for this study. A) Map of the southern region of Kāne‘ohe Bay where the study took place. Orange dots indicate the center of the Outer Lagoon and Inner Lagoon patch reefs. Inset shows the island of O‘ahu, with the red box indicating the southern region of Kāne‘ohe Bay. B) Schematic of coral collection, fragmentation, and reciprocal transplantation. Representative images of the coral species used in this study for C) *M. capitata* and D) *P. compressa*. Images of the experimental setup at the Outer Lagoon reef (E) and the Inner Lagoon reef (F).

In order to characterize the physicochemical dynamics at each reef, temperature, salinity, pH and dissolved oxygen (DO) were measured on the reef benthos at each site (2 m depth) and recorded in 15-minute intervals using SeapHOx sensors (SeaBird Electronics). Prior to deployment, the pH sensors were conditioned in a flow-through seawater aquarium and the DO sensors were calibrated using 100% air-saturated seawater and an anoxic sodium sulfite solution (10 mg ml^-1^). Following deployment, discrete water samples were collected monthly at each site immediately adjacent to the SeapHOx intake at the corresponding time of the instrument sampling. Seawater samples were returned to the lab within 45 minutes of collection and analyzed in triplicate for spectrophotometric pH measurements using *m*-cresol purple as described by SOP 6b (10 cm path length; Dickson et al. 2007). Photosynthetically active radiation (PAR) was measured in 15-minute intervals using Odyssey PAR loggers (three sensors per site) calibrated to a Licor cosine light sensor. Sedimentation rate at each reef was measured using triplicate sediment traps oriented at 90° degrees from the bottom and open at the mouth (Storlazzi et al. 2011), which were collected and redeployed every two weeks. Relative rates of water movement at each reef were measured every ~2-3 weeks (9 times total) using the clod-card dissolution technique (6 cards per reef; (Doty 1971; Jokiel and J Morrissey 1993).

#### Identification of bleaching resistant coral colonies

Bleaching resistant colonies of *Montipora capitata* (Fig. 1C) and *Porites compressa* (Fig. 1D) were tagged during the peak of the 2015 coral bleaching event (Matsuda et al. 2020). One year later, ten individuals of each species at each of the two reefs (40 colonies total), with the exception of two *M. capitata* colonies from the OL reef with unknown bleaching history. *M. capitata* colonies were genotyped using microsatellite markers as described in (Concepcion et al. 2010) in order to confirm that clonemates were avoided. Amplification attempts using primers from (Baums et al. 2012) for *P. compressa* were unsuccessful, but individuals sampled were spaced at least 5 m apart and on average ~20 m apart on the reef to minimize chances of collecting clones.

#### Reciprocal transplant

A portion of each of the 40 coral colonies described above was collected in August 2016 under HIMB Special Activities Permit 2016-69. Each colony was fragmented into 80 small (~4 cm) fragments (Fig. 1B) for use in physiological analyses and were attached to numbered acrylic frag plugs using cyanoacrylate gel. Two larger fragments (~15 cm^2^; Fig. 1B) per colony were attached to numbered plastic grids using underwater epoxy (A-788 Splash Zone Two Part Epoxy) for use in reproduction assays. All coral fragments were mounted on PVC racks (‘coral racks’), which were attached to underwater frames at their native reef (Fig. 1E-F) within 24 hours of collection. Corals were allowed to recover *in situ* for 5 – 7 days (small fragments) or 1 – 4 days (large fragments) prior to initiation of the reciprocal transplant on August 19, 2016 (Fig. 1C). Half of the fragments from each colony were kept at their origin reef (native site), and half were transplanted to the other reef (cross-transplanted; Fig. 1B). To do so, five replicate fragments per parent colony were randomly distributed onto each of four coral racks at each site by divers, for a total of 100 fragments of mixed origin per rack (5 replicate fragments from each of 20 conspecific parents). For the large coral fragments, 10 conspecific fragments were attached to each coral rack. Coral racks were then haphazardly arranged and secured to the frames (Fig. 1E-F). All coral racks were randomly repositioned along the frames within a site every 6 weeks for the first 6 months of the experiment. After that, only the large coral fragments remained, and these were spread out to 5 corals per rack. The racks were rearranged along the frames within each site monthly until the start of spawning (May 2017). The reciprocal transplant resulted in four transplant histories (i.e. origin x destination crosses).

### Thermal Challenge

Six months following the reciprocal transplant, a subset of coral fragments (2 per genet per transplant history; 160 fragments total) were randomly allocated into an ambient or high temperature treatment group for an acute thermal stress experiment. A total of 8 indoor flow-through seawater tanks were randomly assigned as ambient and high temperature treatment tanks, and 20 corals were placed in each tank. Each tank was illuminated by an LED aquarium light (Ecotech Marine XR30w Pro) on a 12:12 h light:dark cycle set to mimic the *in situ* light cycle (peak of ~730 μmol m^-2^ s^-1^). The temperature of each tank was monitored and controlled using a Neptune Apex Aquarium Controller system in combination with a titanium aquarium heater (Finnex TH Series 300w) and a recirculating water pump (Rio 1100+). All tanks started at a daily “ambient” range of 24.5 – 26°C (“day 0”). High temperature tanks were ramped 1°C per day for 6 days, reaching a maximum of 32°C (MMM+4), and held at this temperature profile for the remainder of the experiment (Fig. S5). Ambient temperatures increased over the course of the experiment due to warming weather, but never overlapped with the high temperature treatment. Apex temperature probe measurements were verified using a Traceable Certified Thermometer (VWR). On days 3 and 6 of the experiment corals were randomly shuffled within each tank. Coral skeletal accretion, photosynthesis and respiration rates for each fragment were determined at the beginning and end of the experiment as described below, with incubations at the respective treatment temperatures. Each run included fragments from multiple experimental tanks, and ambient and high temperature treatments were measured in alternating incubations. Photochemical efficiency (Fv/Fm) of the Symbiodiniaceae was assessed daily as described below.

### Coral fitness response

#### Survival and growth

Survival was monitored weekly in the field beginning at 6 weeks post-transplantation and dead fragments were removed. Coral growth was determined for a subset of fragments using two methods: 1) skeletal accretion was determined using the buoyant weight technique (Davies 1989), and 2) linear extension was determined from photographs taken at a fixed distance that included a ruler and the change in the maximum axial length of each fragment as quantified in ImageJ relative to the standard. Initial size measures were taken for 10 fragments per parent per transplant treatment (N=800) immediately following transplantation and again following 3 months (November 2016), at which point half were sacrificially sampled (see below). The remaining half were returned to the field and assessed again after 6 months of transplantation (N=400; February 2017).

#### Reproductive output

Reproductive output of *M. capitata*, a broadcast spawning simultaneous hermaphrodite (Padilla-Gamiño and Gates 2012), was quantified after 9-11 months of transplantation during the spawning season (May – July 2017). This transplantation period encompassed the entire reproductive development of this species (Padilla-Gamiño et al. 2014), from the start of gametogenesis in late summer of 2016 through its culmination during spawning the following year. Corals were collected from the reef three days prior to the new moon, placed into flow-through seawater tanks under natural light (50% shade) at the Hawai‘i Institute of Marine Biology, and monitored in individual containers for five nights beginning on the night of the new moon. Total reproductive output was determined by measuring the cumulative volume of egg-sperm bundles released from each individual across all nights in the months of June, July, and August. Reproductive output was normalized to planar surface area of live coral tissue measured from overhead photographs of each colony using ImageJ. The number of eggs per bundle was determined from five individual bundles per colony as described in (Padilla-Gamiño et al. 2014). Efforts to quantify reproductive output for *P. compressa*, a gonochoric broadcast spawner (Neves 2000), were unsuccessful.

#### Fitness score calculation

The fitness of coral transplants, defined as the relative contribution of individuals to the next generation, is dependent upon their ability to survive and reproduce in their new environment. In colonial organisms like corals, reproductive output is positively correlated with coral size (Hall and Hughes 1996), making growth an important metric for predicting coral fitness. We therefore developed a cumulative metric of coral fitness (i.e. fitness score) that compiled coral survival, growth (skeletal mass), and for *M. capitata* only, reproductive success. The proportion of individual fragments from each genet and history that survived at 6 months was multiplied by their respective growth (represented as the total change in skeletal mass across the 6 months), and for *M. capitata* only, by the proportion of genets from each history that successfully reproduced following transplantation.

#### Local specialization calculation

Local specialization (S) was calculated as the difference in fitness (W) between the home and transplanted environment for each genet, divided by the mean fitness of all corals of that species at that transplant site, regardless of origin (following (Hereford et al. 2009; Kenkel et al. 2015). Positive values indicate native genets perform better than transplanted corals; while negative values indicate genets perform better when not at their native reef (i.e., cross-transplants).

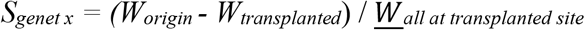

### Coral metabolic traits

#### Oxygen Flux

Photosynthesis and light enhanced dark respiration (LEDR) rates were quantified after 3 and 6 months of transplantation (N=400 per time point: 5 fragments per parent per site; see results for actual totals after mortality). Corals were brought in from the field, placed in flow-through seawater tables, cleaned of any fouling organisms, and assessed within 8-hours of collection. Each coral was placed into a 250 mL respirometry chamber filled with ambient seawater and sealed. One control chamber (seawater only) was run alongside each round of coral incubations. Corals were maintained at constant ambient temperature (T3: 24°C; T6: 23.5°C) using a water jacket, and the water within each chamber was mixed with a magnetic stir bar. Temperature and dissolved oxygen concentrations were measured simultaneously using a Pt100 temperature probe and PSt7 oxygen optode (Presens), respectively, which were inserted through ports in the lid of each chamber. Data were recorded once per second via an OXY-10 ST (Presens). Optodes were calibrated with a 0% oxygen solution (10 mg ml^-1^ NaSO_3_) and 100% air saturated seawater. Rates of photosynthesis were measured under 700 μmol m^-2^ s^-1^ light (equivalent to midday daytime light levels on the reefs; Fig. S1) until a steady rate was observed for at least 10 minutes for all corals. At this point, the lights were turned off and light enhanced dark respiration (LEDR) rates were determined until a steady slope was obtained for at least 10 minutes. The amount of oxygen released or consumed over time was calculated by multiplying by the oxygen concentrations with the volume of water (volume of the chamber less the volume of the coral). Coral volumes were calculated from the mass of each coral: Volume = mass/Density; skeletal density was empirically determined (see Supplemental Methods). The rate of oxygen evolution was determined from a linear regression and rates were corrected for blank chamber rates and normalized to coral surface area.

#### Photochemical activity

Dark adapted photochemical efficiency (Fv/Fm) was assessed after 1.5, 3, and 4.5 months post-transplantation using PAM fluorometry. Corals (10 fragments per parent per site; 800 total) were brought in from the field in the late afternoon and kept in flow-through seawater tables under natural light (50% shade). Corals were dark-adapted for at least 30 minutes following sunset, and photochemical efficiency of photosystem II (Fv/Fm) was measured using a Dive-PAM (Walz GmBH). Corals were returned to the field the following morning.

#### Heterotrophic activity

Coral heterotrophic activity was quantified for a subset of the above corals (one per parent: 10 per history per species; N=80) at 3 and 6-months post-transplantation (modified from Towle et al. 2015). Coral fragments were isolated in an indoor tank in 1 μm filtered seawater (FSW) for a 6-hour heterotrophic deprivation period the same day of collection from the field. One hour after sunset, corals were placed in 220 mL chambers in 1 μm FSW with magnetic stirring. After a 30-minute acclimation period, a natural assemblage of plankton (including copepods, zoea, and phytoplankton) were added to each chamber (3,000 plankters L^-1^) for 60-minutes after sunset (Levas et al. 2016). Triplicate 1 mL water samples were collected from each chamber after 0, 30, and 60-minutes, and the number of plankters was immediately counted under a dissecting microscope. Chambers without corals were used to calculate background rates of prey depletion in the absence of coral feeding (N=4). Feeding rate was calculated as the decrease in total number of plankters corrected for blank chamber rates and normalized to coral surface area.

#### Biomass

Coral fragments were flash-frozen in liquid nitrogen following assessment of autotrophic or heterotrophic activity, and stored at −80°C. Tissue was removed from the skeleton using an airbrush with 0.2 μm filtered seawater (FSW). The resulting homogenate was dried at 60°C to constant weight, then burned at 400°C for 4 hours. Biomass, defined here as ash free dry weight (AFDW), was calculated as the difference between the dry weight and the ash weight.

#### Lipids

Coral tissue total lipid content was quantified gravimetrically (modified from Wall et al. 2019) by extracting lipids from coral tissue slurries in 2:1 chloroform:methanol with 0.88% KCl and 100% chloroform washing. Lipids were filtered onto pre-combusted (450°C, 6 h) GF/F filters, dried at 60°C (10-12 h) followed by combustion at 450°C (6 h). The remaining slurry (non-lipid fraction) was used to quantify biomass by AFDW as described above. Lipid and biomass fractions were weighed before and after combustion to the nearest 0.0001 g. Lipids were calculated as a proportion of total tissue biomass (lipids (g) + biomass (g)).

#### Normalization

Coral skeletons were soaked in 10% bleach overnight to remove organic matter, dried to constant weight at 60°C, and skeletal surface area was determined by the wax dipping method (Veal et al. 2010). These values were used for normalization of autotrophic and heterotrophic activity.

### Statistical analysis

Statistical analysis of coral response to transplantation were conducted in R Statistical Programming (R Core Team 2017). Univariate analyses were performed using a linear mixed effect model approach in the *lme4* (Bates et al. 2015) and *glmmADMB* (Skaug et al. 2016) packages. Proportion survivorship was analyzed by a beta regression in the *betareg* package (Cribari-Neto and Zeileis 2010). Fixed effects (origin, destination, species) and random intercepts (genotype, rack) of models for each coral response are described in detail in Table S1. For all models, assumption of residual normality was assessed using quantile-quantile plots and homogeneity of variance of residuals was assessed using the Levene’s test in the *car* package (Fox and Weisberg 2018). Transformations were determined by AIC model selection. Heterotrophic feeding and buoyant weight data were square root transformed and biomass and dark-adapted yield data were log transformed to meet assumptions of analyses, but data are plotted as untransformed values. Significance testing was completed using Type III ANOVA sum of squares tests in the *lmerTest* package (Kuznetsova et al. 2017) and *car* packages (Fox and Weisberg 2018). Principal component analysis (PCA) was used to determine the percent variance explained by seven physiological variables (biomass, calcification, linear extension, gross photosynthetic rate, LEDR, P:R, and survival) in the separation of the transplant groups, using data from all genets, and to quantify the extent of plasticity exhibited by each origin population. PCA was conducted on the scaled and centered data using the *prcomp* function in the Vegan package (Oksanen et al. 2007). Given that the first two PCs explained the majority of the variance, phenotypic plasticity of each genet was calculated as the PCA distance between that genet’s native vs. cross-transplanted phenotype in two-dimensional trait space (i.e. PC1 vs. PC2), which accounts for correlations among traits (as in (Abbott et al. 2018) and is reported as plasticity. Differences in plasticity were tested using a two-way ANOVA with the factors of Species and Origin.

## Results

### Distinct physicochemical dynamics characterized each reef

Temperature dynamics did not differ significantly between the two patch reefs, with mean temperatures of 25.23 ± 1.55°C at the Outer Lagoon reef and 25.14 ± 1.56°C at the Inner Lagoon reef across the six-month transplantation period, and a corresponding mean diel range of 0.50 ± 0.16°C vs. 0.52 ± 0.16°C, respectively (Fig. 2A-B; Table 1). Mean pH and dissolved oxygen (DO) were also similar between the two reefs (Fig. 2, Table 1), as were light levels (Fig. S1; Table 1). Diel fluctuations in pH and DO were significantly greater at the Outer Lagoon than the Inner Lagoon reef (Fig. 2), with the daily pH amplitude 2.92-fold higher at the Outer Lagoon reef than the Inner Lagoon reef (0.111 pH units day^-1^ vs. 0.038 pH units day^-1^, respectively) and 2.68-fold higher for DO at the Outer Lagoon reef than the Inner Lagoon reef (73.9 μM day^-1^ vs. 27.6 μM day^-1^, respectively; Table 1). Sedimentation rates were 8.27-fold higher at the Outer Lagoon (0.324 ± 0.066 g day^-1^) than the Inner Lagoon reef (0.039 ± 0.003 g day^-1^; Wilcoxon rank sum test, p<0.0001; Table 1). Relative flow rates were also ~2-fold higher at the Outer Lagoon reef (t-test, p<0.0001; Table 1). In contrast, diel fluctuations in salinity were lower at the Outer Lagoon than the Inner Lagoon reef (Fig. 2E-F, Table 1).

**Figure 2.**
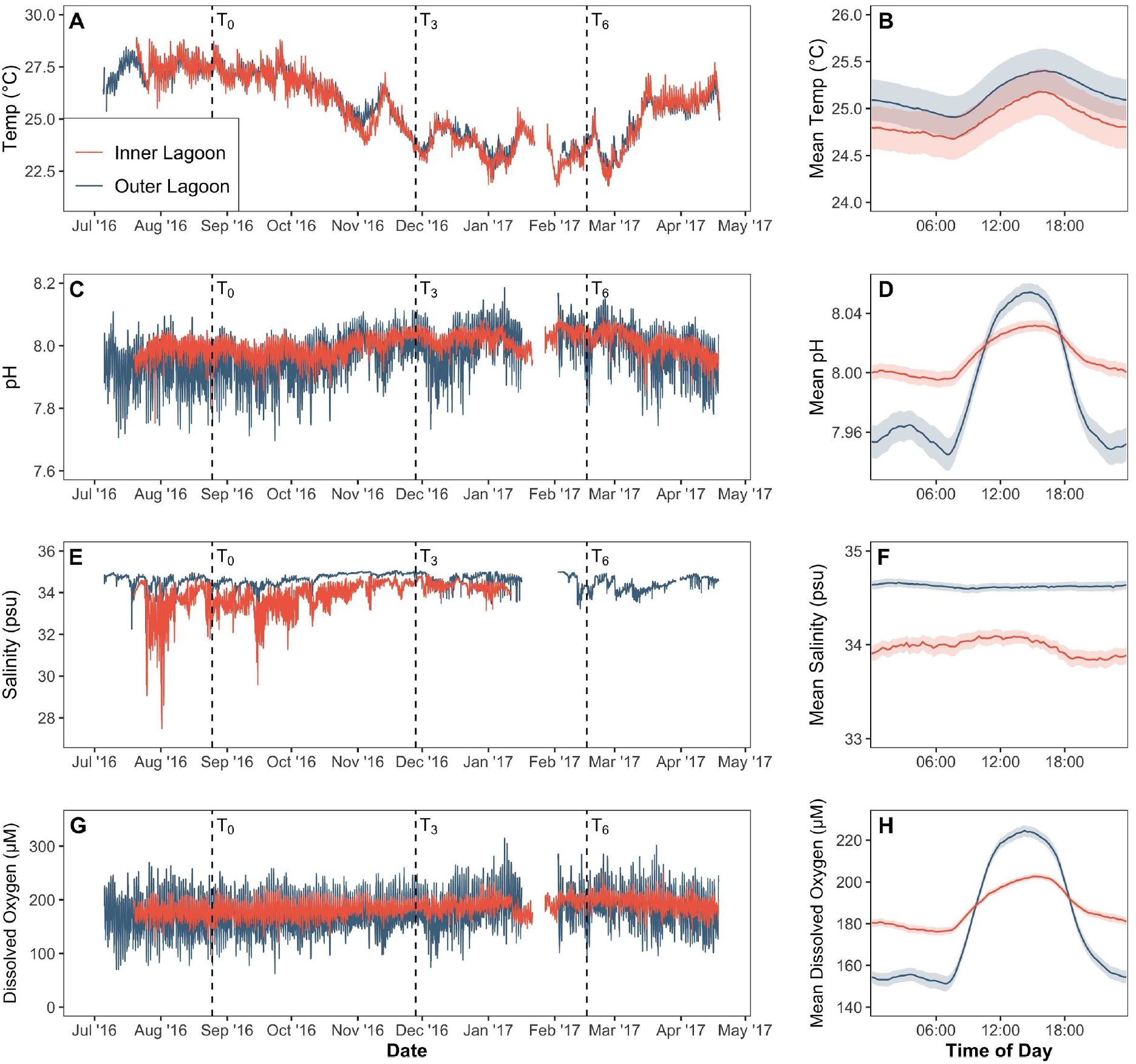
Characterization of seawater physicochemical dynamics above the reef benthos at the Inner Lagoon (orange) versus Outer Lagoon (blue) reefs. Seawater temperature time series (A) and mean diel temperature cycle (B). Seawater pH time series (C) and mean diel pH cycle (D). Salinity time series (E) and mean diel salinity cycle (F). Seawater dissolved oxygen (DO) content time series (G) and mean diel DO cycle (H). Vertical dashed lines indicate the initiation of the transplant (T_0_) followed by sampling time points after 3 months (T_3_) and 6 months (T_6_) of transplantation. The mean diel cycles are shown with shading indicating ± 95% confidence interval.

**Table 1.** Mean values and daily amplitude (where applicable) for environmental parameters at the Inner and Outer Lagoon reefs. Asterisks denote significant differences (p<0.05).

| Mean | Inner Lagoon | Outer Lagoon | Ratio (OL:IL) |
| --- | --- | --- | --- |
| Temperature (°C) | 25.137 ± 1.555 | 25.225 ± 1.547 | 1.00 |
| pH | 8.007 ± 0.028 | 7.983 ± 0.049 | **1.00\*** |
| Dissolved oxygen (μM) | 185.8 ± 8.3 | 177.9 ± 13.2 | **0.96\*** |
| Salinity (psu) | 33.98 ± 0.51 | 34.66 ± 0.27 | **1.02\*** |
| Daily light integral (mol m<sup>-2</sup> d<sup>-1</sup>) | 11.3 ± 5.24 | 12.3 ± 5.53 | 1.09 |
| Sedimentation rate (g day<sup>-1</sup>) | 0.039 ± 0.003 | 0.324 ± 0.066 | **8.27\*** |
| Flow rate (% dissolution h<sup>-1</sup>) | 1.22 ± 0.03 | 2.35 ± 0.03 | **1.92\*** |
| Daily Amplitude | Inner Lagoon | Outer Lagoon | Ratio (OL:IL) |
| Temperature (°C) | 0.515 ± 0.161 | 0.499 ± 0.161 | 0.97 |
| pH | 0.038 ± 0.013 | 0.111 ± 0.041 | **2.92\*** |
| Dissolved oxygen (μM) | 27.6 ± 9.5 | 73.9 ± 28.7 | **2.68\*** |
| Salinity (psu) | 0.257 ± 0.08 | 0.06 ± 0.014 | **0.23\*** |
| Light (μmol m<sup>-2</sup> s<sup>-1</sup>) | 590.5 ± 21.1 | 597.0 ± 29.2 | 1.01 |

### Coral acute heat stress response was unaffected by transplantation

At the initiation of the acute heat stress experiment (maximum daily temperature of 27°C across all treatments; Fig. S8), there were no significant differences in performance metrics (i.e., LEDR, Fv/Fm, and gross photosynthesis) between species, treatments, origins, or destinations (Fig. S9). At the end of the 10-day heat stress, the heat treatment reached a daily maximum of 32°C (maximum monthly mean [MMM] +4°C), while the ambient treatment reached a daily maximum of 28°C (Fig. S8). At this point, linear mixed models (Table S21) indicated that treatment was a significant factor across all parameters examined, with corals in the heat treatment exhibiting declines in photochemical yield (Figure 3A; Table S22), metabolic rates (Figure 3B,C; Table S23), and calcification rates (Figure 3D, Table S24). For corals in the heat treatment, origin was not a significant factor for any of the metrics examined (Tables S22-24), indicating that there was no change in bleaching resistance six months following transplantation to a novel environment. This was true for cross-transplanted corals relative to their performance at their native reef (i.e. between destinations), and for cross-transplanted corals relative to native corals within each reef (i.e. within destinations; Figure 3). Species was also a significant factor for photochemical yield (Table S22), photosynthesis (Table S23), and calcification rates (Table S24), but not respiration (LEDR) rates (Table S23). Overall, *P. compressa* showed the greatest declines in performance metrics in response to heat stress, with declines in photochemical yield, metabolism, and calcification exceeding those of *M. capitata* (Fig. 3A-D). In contrast to the pattern seen in the field, coral performance was not influenced by destination at the end of the acute heat stress experiment (Table S22-24). There was however a significant 3-way interaction between destination, species and treatment for calcification (Table S24), driven by lower calcification of Outer Lagoon natives of *P. compressa* at ambient conditions relative to cross-transplants under ambient conditions acclimatized to that site (Fig. 3D).

**Figure 3.**
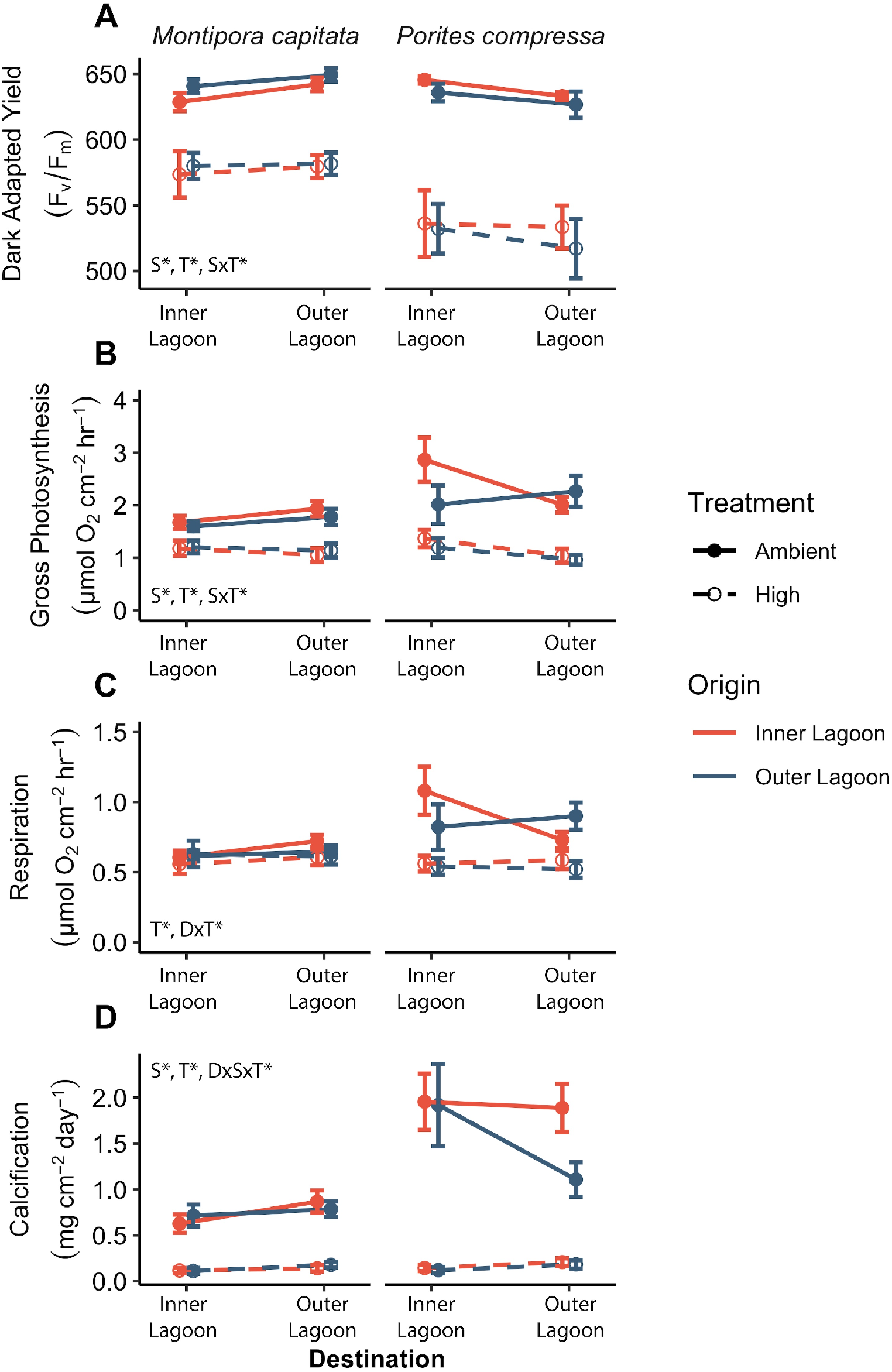
Coral performance following acute thermal stress (high temperature; 32°C) vs. controls (ambient temperature; 27-28°C). A) Photosynthetic efficiency (dark adapted yield; Fv/Fm), B) gross photosynthesis rates, C) respiration rates, and D) calcification rates. N=8-10; error bars indicate SEM.

### Coral fitness differed between reefs but no evidence of site specialization

Overall, corals at the Outer Lagoon reef showed significantly higher fitness scores than corals at the Inner Lagoon reef. There were, however, no differences in fitness between native and cross-transplanted corals at the Outer Lagoon (Figure 4A,B, Table S19-20). In contrast, at the Inner Lagoon reef, cross-transplants of *M. capitata* displayed higher fitness than native corals, but only when accounting for differences in reproductive success (Figure 4A, Table 19-20). Absence of reproductive data from the *P. compressa* fitness score calculation implicitly assumes 100% reproductive success, so these values could only decrease with the inclusion of additional data. Sequencing of microsatellite markers confirmed that the *M. capitata* colonies sampled were not clones (Table S25), however genotyping of *P. compressa* was unsuccessful. Despite lack of molecular confirmation for *P. compressa*, it is unlikely that the individuals used in this study were clones because of distance apart on the reef and low levels of clonality within the lagoon (Locatelli and Drew, n.d.). Local specialization of each genet was calculated by considering the relative fitness of that genet in its native vs. cross-transplanted environment. In general, corals exhibited positive specialization values at the Outer Lagoon reef, whereas corals native to the Inner Lagoon reef showed negative specialization values (Figure 4B), indicating corals performed better at the Outer Lagoon reef even when it was not their native environment. The only exceptions were one genet of *M. capitata* native to the Outer Lagoon, which had a negative local specialization score, indicating it had higher fitness when cross-transplanted to the Inner Lagoon, and one genet of *P. compressa* native to the Inner Lagoon that had a positive local specialization score and was thus the only genet of either species native to the Inner Lagoon that had higher fitness at its native reef (Figure 4B).

**Figure 4.**
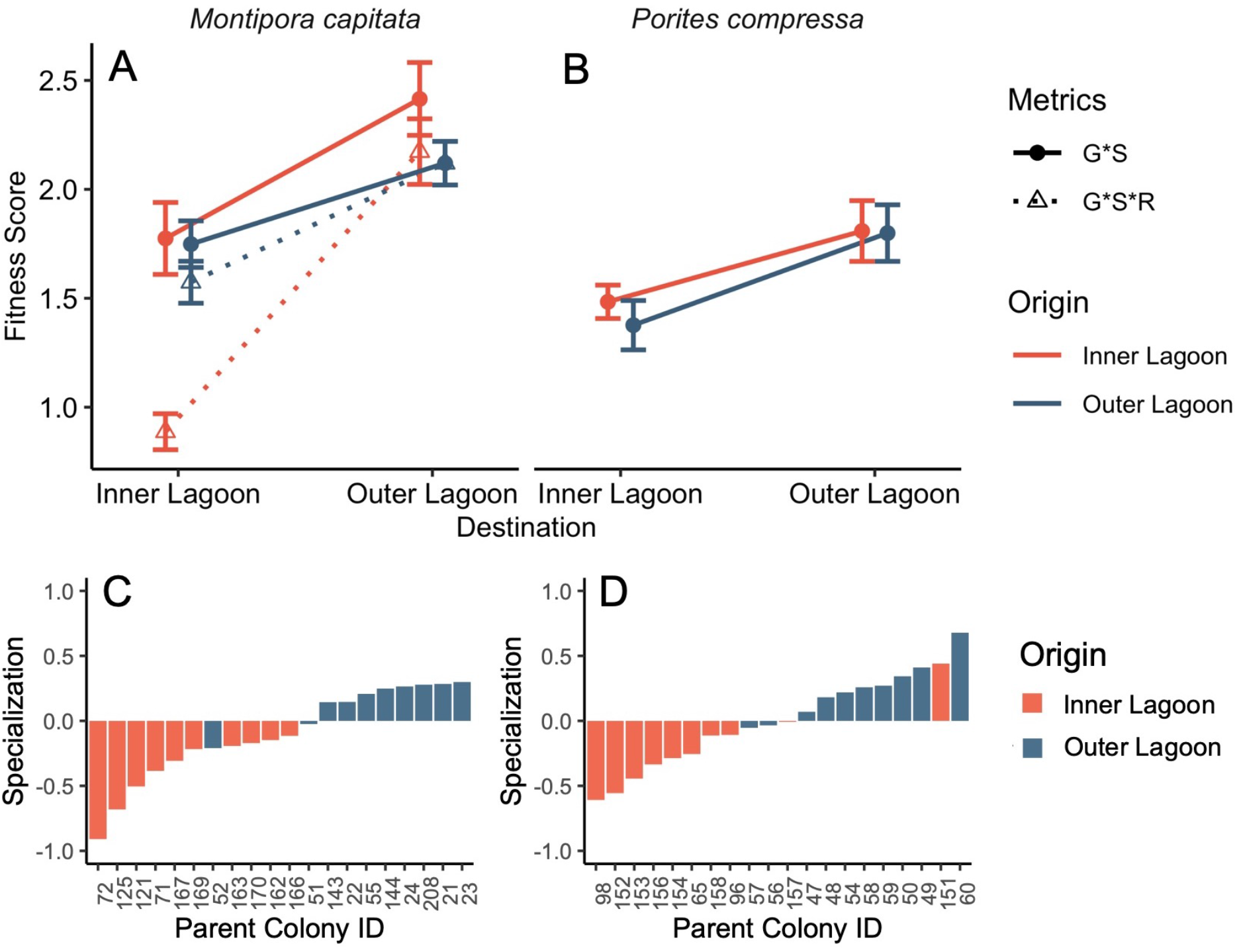
Fitness score for A) *Montipora capitata* and B) *Porites compressa*. Fitness score is a product of survival (S), net growth (G), and for *M. capitata*, reproductive success (R). N=10; error bars indicate SEM. Magnitude of local specialization for each genet of C) *M. capitata* and D) *P. compressa*. Local specialization values are defined as the difference in fitness score (G*S only) of a genet at its origin and destination reef, divided by the mean fitness score of all conspecifics at the destination reef. Positive values indicate local site specialization; negative values indicate destination reef favorable.

### Corals exhibited high phenotypic plasticity in response to novel reef environments

Significant differences in coral phenotypes were observed between destinations and species after both three and six months following transplantation (p=0.001; PERMANOVA; Fig. 5A,B). There was also a significant interaction between species and destination at both time points (p=0.003 for T3 and p=0.001 for T6; PERMANOVA). In contrast, origin was not a significant factor at either time point, indicating that both species had acclimatized to their destination reef as early as three months following transplantation. Genotype plasticity, quantified as the PC distance between each genet’s native vs. cross-transplanted phenotype, did not differ between the two origin populations for either species. Plasticity did differ between species, with *P. compressa* exhibiting higher phenotypic plasticity than *M. capitata* at both T3 (p<0.0004) and T6 (p=0.017) (Fig. 5C, 5F). The traits contributing most strongly to differences between destinations included metabolic rates and growth, which were higher at the Outer Lagoon reef after three months, whereas biomass and survival were higher at the Inner Lagoon reef (Fig. 5A). After six months post-transplantation, all traits were higher at the Outer Lagoon reef. Comparing species, *P. compressa* had higher biomass, P, and R, while *M. capitata* exhibited higher survival, growth, and P:R (Fig. 5D).

**Figure 5.**
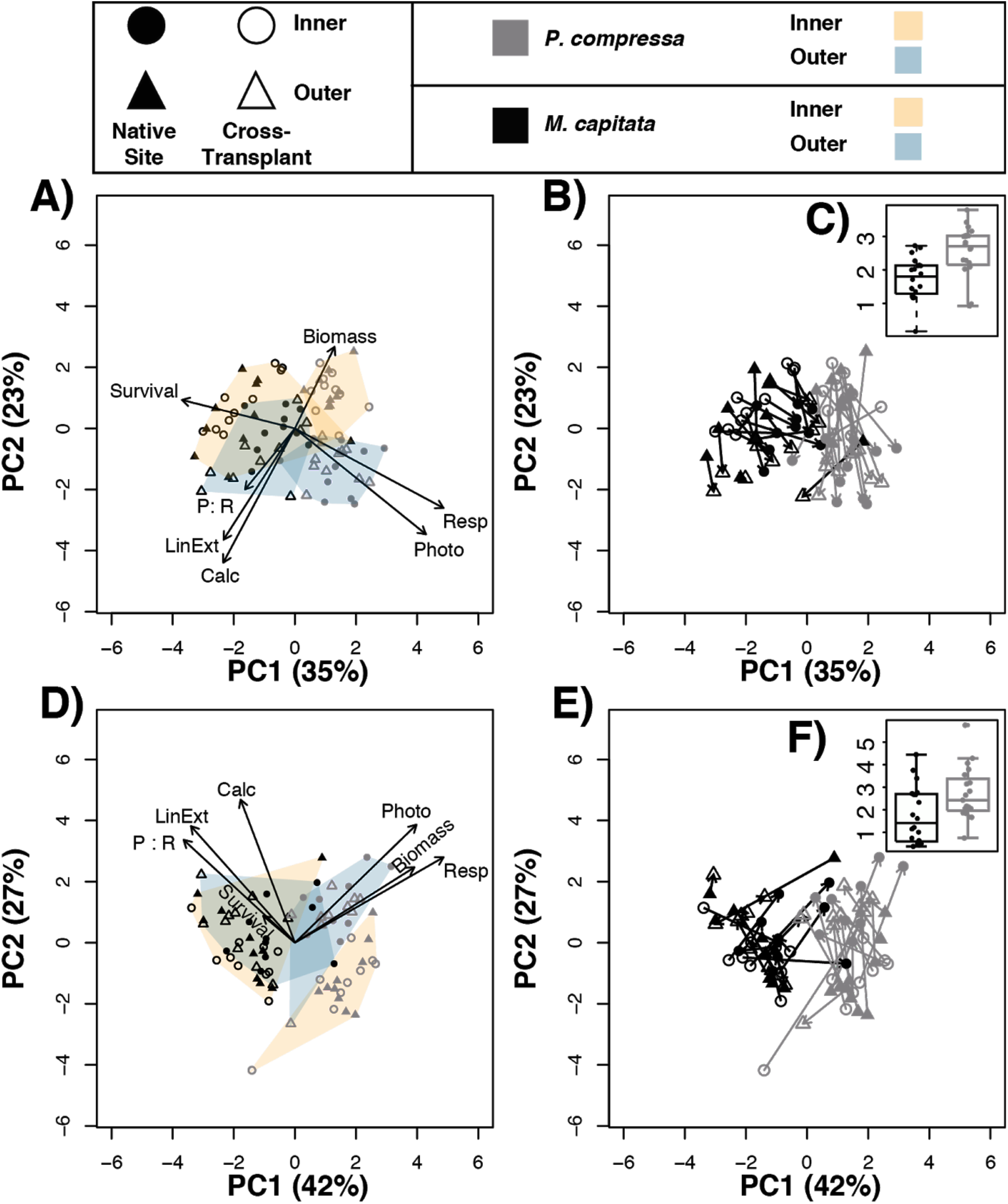
Principal component analysis of coral performance following three (A-C) and six months (D-F) of transplantation. Polygons outline the ordination groups, with *Porites* in purples and *Montipora* in greens, whereas vectors in (A) and (D) indicate the loadings of the phenotypic variables to the PCs, with length of arrow signifying strength of loading. Plasticity, calculated as the distance in principal component space between each genet’s native (filled symbols) vs. cross-transplanted phenotype (open symbols) are indicated by lines in (B) and (E). The boxplots and data points for plasticity values of each species are shown in (C) and (F).

### Physiological responses following transplantation

#### Metabolic traits

Both species had greater biomass at the Outer Lagoon reef than conspecifics at the Inner Lagoon reef after three months, and *P. compressa* had greater biomass than *M. capitata* across both reefs regardless of origin (Figure 6A, Table S2-3). Lipid content also differed between species and destinations but not origin, with corals at the Outer Lagoon having lower lipid content and *P. compressa* tissues containing lower proportions of lipids than *M. capitata* (Figure S3, Table S4-5). All corals consumed plankton through heterotrophic feeding activity, depleting the prey population at a rate ranging from 7.5 to 27 plankters cm ^-2^ h^-1^. *P. compressa* also had higher feeding rates at the Inner Lagoon reef relative to the Outer Lagoon reef at six months (Figure S4B); *M. capitata* feeding rates were also higher at the Inner Lagoon reef, but only at the three-month time point (Figure S4A, Table S6-7). *P. compressa* had higher feeding rates than *M. capitata* after six, but not three months of transplantation (Figure S4A-B).

**Figure 6.**
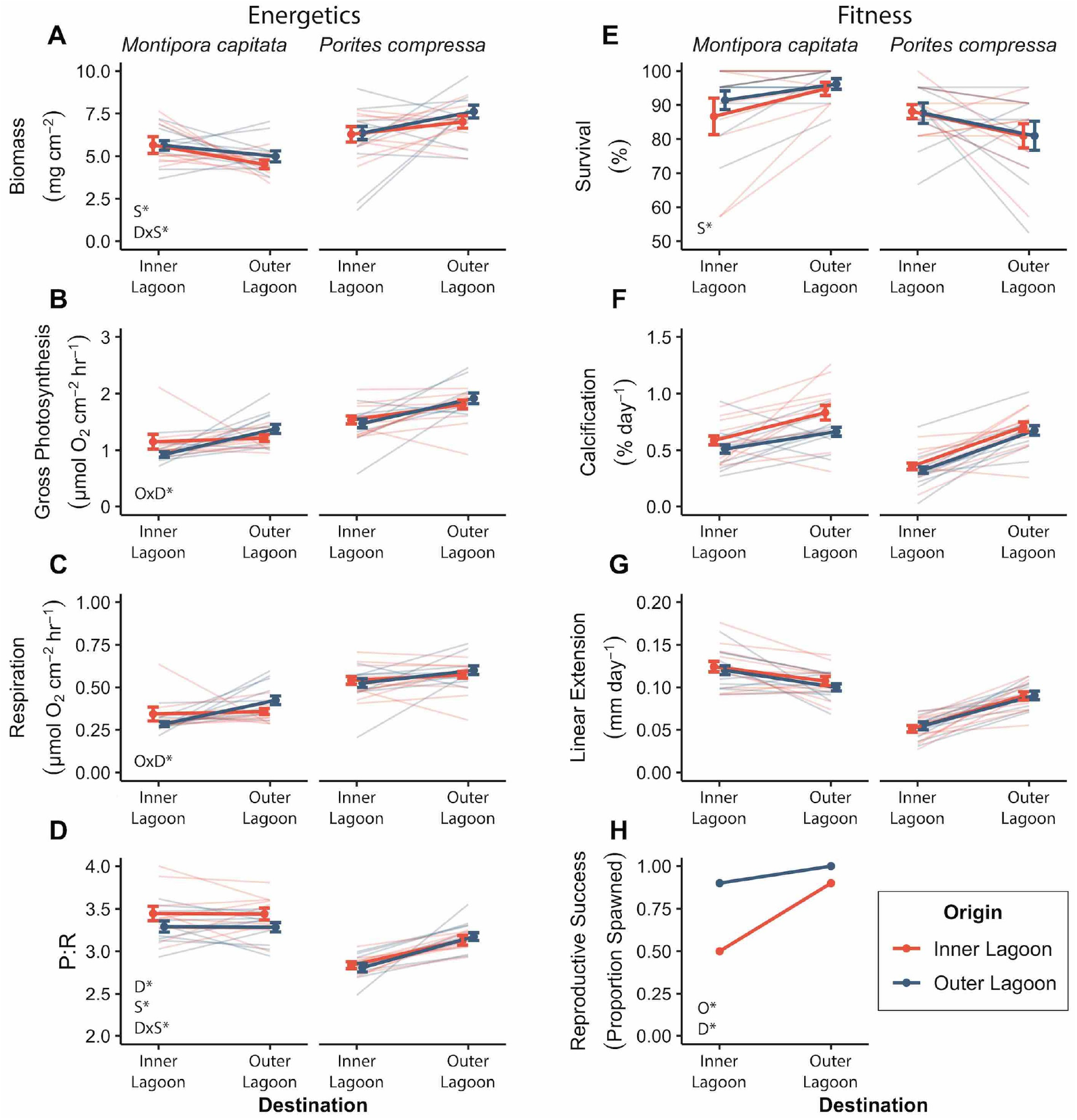
The effects of origin site and transplant site on coral energetics (left) and fitness (right) traits after 6 months following reciprocal transplantation of *Montipora capitata* and *Porites compressa* between an Inner and Outer Lagoon reefs. A) Biomass; B) Gross photosynthesis; C) Respiration; D) Photosynthesis to respiration ratio (P:R); E) Survival; F) Calcification; G) Linear extension; and H) Reproductive success. Bold lines indicate the mean of all genets (N=10) ± standard error of the mean (SEM); thin lines indicate mean response of each genets (N=5 ramets).

Autotrophic activity was measured by assessing photochemical efficiency and photosynthetic rates of the coral ramets. Photochemical efficiency (Fv/Fm) did not differ between origins or destinations, but was higher for *M. capitata* than *P. compressa* after 1.5 months of transplantation (Figure S5A, Table S8-9). After three months, both species and destination were significant factors, but at 4.5 months none of the factors were significant (Figure S5B-C, Table S8-9). Gross photosynthesis rates also differed between destinations and species after three months of transplantation, where they were higher overall at the Outer Lagoon reef and higher for *P. compressa* than *M. capitata* within each reef (Figure S2B, Table S10-11). In contrast, six months after transplantation there was a significant interaction between origin and destination, as corals of both species originating from the Outer Lagoon reef showed a decline in photosynthesis rates at the Inner Lagoon reef relative to the Outer Lagoon reef whereas corals originating from the Inner Lagoon reef showed no difference in photosynthesis rates between their native Inner Lagoon vs. the Outer lagoon reef (Figure 6B, Table S10-11). Respiration rates also differed between species and destinations after three months of transplantation, and similar to photosynthesis were higher at the Outer Lagoon reef than the Inner Lagoon reef and higher for *P. compressa* than *M. capitata* (Figure S2C, Table S10,12). After six months of transplantation, similar again to photosynthesis rates, species and destination were no longer significant factors for respiration, but there was a significant interaction between origin and destination (Figure 6C, Table S10). The relative ratios of photosynthesis to respiration (P:R) were not different between species, destination, or origin after three months of transplantation (Figure S2D, Table S10). In contrast, after six months of transplantation, species and destination were significant factors, and there was a significant interaction between the two where P:R was higher at the Outer Lagoon for *P. compressa* but lower at the Outer Lagoon for *M. capitata* relative to the Inner Lagoon (Figure 6D, Table S10). In addition, *M. capitata* had higher P:R than *P. compressa* at the Inner Lagoon, which was driven by the relatively high respiration rates of *P. compressa* vs. *M. capitata* at that site that balanced out the higher photosynthesis rates of *P. compressa* at that site (Figure 6D, Table S13).

#### Survival, growth and reproduction

Species was the only significant factor affecting survival at both time points, which was higher for *M. capitata* than *P. compressa* at both reefs (Figure 6E, Figure S2E, Tables S14-15). Destination was the only significant factor affecting calcification, which after three months was higher for both species at the Outer Lagoon reef relative to the Inner Lagoon reef regardless of origin (Figure S2F, Table S16-17). However, these differences were no longer significant after six months (Figure 6F). Linear extension did not differ between species, origin or destination at either time point (Figure 6G, Figure S2G, Tables S16,S18). Reproductive success, quantified as the proportion of *M. capitata* genets from each transplant group that successfully spawned, was significantly affected by both origin and destination (Table S14). Reproductive success of *M. capitata* at the Outer Lagoon reef was higher than at the Inner Lagoon reef and did not differ between origins (Figure 6H), whereas at the Inner Lagoon reef native *M. capitata* had lower reproductive success (50%) than cross-transplants (90%; Figure 6H). This is the only metric examined in this study where origin alone was a significant factor. Reproductive output was not significantly different between origin or destination (mean 0.018 ± 0.019 mL cm^-2^ live tissue; Figure S6). Cross-transplants at the Inner Lagoon reef (i.e. introduced from the Outer Lagoon reef) showed a strong positive relationship between growth and reproduction (p<0.005, r^2^=0.700; Figure S7). In contrast, natives at the Inner Lagoon showed no relationship between growth and reproduction. Likewise, at the Outer Lagoon reef there was no relationship between growth and reproduction for either native or cross-transplanted corals, indicating that there were no negative tradeoffs between growth and reproduction.

## Discussion

### Fitness consequences of coral acclimatization to novel environments

Science-based restoration is key to the success of reefs restored through human intervention. Here, we show transplantation of bleaching-resistant corals to a novel environment *in situ* did not alter their heat stress response, despite transplants exhibiting high levels of phenotypic plasticity for other traits. Because bleaching-resistant individuals have lower mortality (Matsuda et al. 2020) and higher reproductive success (Fisch et al. 2019; Ward et al. 2000; Howells et al. 2016) than bleaching-sensitive conspecifics following a bleaching event, they have a clear selective advantage during and in the years following these events. Harnessing these natural advantages by propagating bleaching-resistant individuals is a promising approach to increase the abundance of corals with these traits, and could potentially increase the bleaching resistance of a reef using native (i.e. endemic, local) coral stocks. Furthermore, because bleaching resistance in these species can persist through multiple bleaching events (e.g. 2014 vs. 2015, Ritson-Williams 2020), these individuals are likely to retain their thermal tolerance across longer time periods than the current study (6-11 months). *Montipora capitata* and *P. compressa* represent divergent lineages of two globally distributed coral genera, suggesting these patterns could be common to other species. Taken together, these results indicate that bleaching resistance is both consistent through time and unaffected by transplantation to a novel environment, and is thus a useful trait for selecting corals for propagation and outplanting to enhance resistance of coral populations to climate change.

Prior to implementing selective propagation for desired traits, consequences on fitness must be assessed and understood. For instance, bleaching-resistant individuals introduced to a new site during non-bleaching years should persist without substantially lowering the fitness of the recipient population even in the absence of a heatwave. Encouragingly, our results demonstrate that there were no negative effects on fitness of recipient populations when new genets were introduced, as the recipient population’s fitness either increased (at the less-favorable site) or stayed the same (at the favorable site) following introduction of bleaching resistant corals during a non-bleaching year. The duration of the observed enhanced fitness, which lasted at least 11 months, remains unknown, as they could be the result of a temporary carryover of the energetic benefits of having originated from a more favorable reef environment. However, even if this carryover is transient, the long-term fitness effects of their introduction are likely net positive due to the transplants’ bleaching resistance, as discussed above. These results are a necessary first step to validate this trait-guided approach to reef restoration. The next important step is to determine if these traits can persist and spread throughout a recipient population, which requires traits to be both heritable and introduced in sufficient abundance. Initial studies indicate that stress-resistant corals must be introduced in numbers equivalent to at least 2-5% of the population per year for several decades in order to achieve adaptive gains in heat tolerance that can keep pace with climate change (Bay et al. 2017). As such, work is needed to scale up these approaches if they are to have a meaningful impact on coral reef resistance to ocean warming.

### Acclimatization via physiological plasticity did not lead to negative tradeoffs

Acclimatization via plasticity can lead to negative trade-offs, where improvements or maintenance of one trait (e.g. growth) come at the expense of another (e.g. reproduction), making identification of possible trade-offs important both for understanding coral biology and for informing trait-guided restoration. Here, transplantation revealed high levels of phenotypic plasticity across a range of traits including metabolism, feeding, growth, and reproduction. Regardless of the directional change, acclimatization via plasticity did not result in negative tradeoffs for any of the traits examined. Critically, plasticity in growth and metabolism did not alter coral heat stress responses for either species, even though *P. compressa* exhibited greater plasticity than *M. capitata*. Also important was the lack of tradeoffs between growth, reproduction, and survival. For example, *M. capitata* genets that increased growth rates following transplantation also exhibited increased reproductive success with no differences in survivorship. Conversely, genets with the largest declines in growth following transplantation also had lower reproductive success, indicating that a lack of investment in growth was not compensated for by increased investments in reproduction. While this study cannot speak to effects of transplantation on *P. compressa* reproductive tradeoffs, it appears that corals at a favorable site performed better across all performance metrics, with none coming at the cost of another. These results are consistent with data from other reef systems that found absence of trade-offs with bleaching and reproduction (Lenz 2020) and resistance to multiple stressors (Wright et al. 2019), and holds promise that these bleaching-resistant genets may also withstand additional stressors.

The magnitude of phenotypic plasticity was greater for *P. compressa* than *M. capitata*, aligning with recent work showing that *M. capitata* calcification is less sensitive than *P. compressa* to differences in environmental conditions (Barnhill et al. 2020). The differences observed here may be due to differential responses to the lower flow regime at the Inner Lagoon reef, which can lead to greater accumulation of metabolic wastes immediately surrounding the coral surface and could have depressed growth and metabolism (Mass et al. 2010). *Porites compressa* may be more sensitive to metabolic inhibition under low flow because of its higher biomass (i.e. thicker tissue) and higher respiration rates than *M. capitata*. Alternatively or in addition, differences in plasticity could have been due in part to morphological plasticity (Todd 2008), as *M. capitata* grew longer, thinner branches at the Inner Lagoon reef than the Outer Lagoon reef, which could thin boundary layers and increase diffusive exchange between the tissues and surrounding seawater (Patterson 1992). Finally, differences in mixotrophy could help explain these responses, as heterotrophic feeding can sustain metabolic demands during environmental changes (Fox et al. 2018), and could have influenced metabolic rates and growth.

### Genotype-environment effects

Both species exhibited variation between genets in the magnitude and direction of their physiological response to each environment. Despite consistently higher mean coral performance at the Outer Lagoon reef, in many cases these differences in performance between the two reefs were not significant due to a strong genotype-environment (GxE) effect, where some individual genets showed higher performance at the Inner Lagoon for some traits. Such differences driven by GxE effects within this cohort of bleaching resistant corals create critical challenges in the selection of individuals with desired climate change resistant traits for coral reef restoration efforts, while also accounting for future performance in other important traits. Growth in particular showed a strong GxE effect, and our findings align with recent work cautioning against using growth alone as a predictive trait for future coral performance as it can vary between seasons (Edmunds and Putnam 2020; Edmunds 2017) and heat tolerance in a stressful environment does not correspond with rapid growth in a less stressful environment (Bay and Palumbi 2017). In summary, this study indicates that bleaching resistance is not plastic in these species and is thus an informative trait for predicting future coral performance. Furthermore, these results argue for using genetically diverse ‘planting stock’ to account for the wide range of expressed phenotypes in different reef environments (Drury et al. 2017).

### Biologically guided strategies for coral reef restoration

There is mounting evidence that ‘natural’ dispersal of heat tolerant genets and the generation times required for adaptation to increase heat tolerance of coral populations cannot keep pace with ocean warming (Quigley et al. 2019). The rapid decline of coral reef habitats accentuates the need for human interventions in management and restoration, such as coral propagation and outplanting. Here we show that bleaching resistance in corals was maintained following introduction to novel environments. Bleaching resistance has shown high heritability in other species (Yetsko et al. 2020; Quigley et al. 2020), and consistent relative ranking of individuals in a common garden setting (Morikawa and Palumbi 2019). While more work is needed to determine how well bleaching resistance persists across generations, these results favor active restoration for promoting climate-ready reefs as the sources of climate change become properly managed. Bleaching resistant genets can be identified either during bleaching events (as in this study) or standardized acute heat stress response assays (Voolstra et al. 2020). Both approaches can facilitate rapid identification of bleaching resistant genets. These efforts could also focus on sites with characteristics known to foster stress tolerant populations (Palumbi et al. 2014). For example, local reefs with high diel variation in temperature, shallow bays with lower flow and thus higher mean temperatures often harbor heat resistant individuals. One such example of this is Kāne‘ohe Bay, Hawai‘i, where corals have higher heat and acidification tolerances than conspecifics from neighboring reefs (Coles et al. 2018; Jury and Toonen 2019; Schoepf et al. 2017) due to the higher mean temps and lower pH of the bay relative to nearby reefs (Drupp et al. 2011). Additional traits are also important when selecting individuals for restoration (e.g. ocean acidification tolerance, disease resistance and genetic diversity; (Muller et al. 2018) plasticity of many of these traits are not well described. Encouragingly, relative growth during acidification stress is consistent in several coral species (Jury et al. 2019), and thus along with bleaching resistance may be a useful selection marker for promoting climate-ready reefs via active restoration.

Site selection for nurseries and outplanting is also an important consideration to maximize restoration success. Here we found that the reef with the greatest water flow, diel physicochemical variation, and distance from land resulted in higher coral growth and fitness than the other reef. High flow and high variability have also been found to promote coral performance in other reef systems (Sully et al. 2019; Safaie et al. 2018) and can mitigate bleaching responses (Page et al. 2019), indicating that these may be generalizable characteristics of high fitness and bleaching resistant coral reefs for many species, though not all (Klepac and Barshis 2020). Our results highlight the importance of selecting sites that promote high coral fitness for nurseries and outplant sites, as this could accelerate the successful establishment of corals following outplanting. Furthermore, an *in situ* nursery site that promoted faster growth would provide obvious logistical benefits, as it would lead to shorter residence time in the nursery and yield greater coral biomass for outplanting. Assisted gene flow using climate-ready genets could complement traditional conservation measures such as marine protected areas (MPAs), which could provide favorable habitat for stress-resistant outplants, and in coordination with less directed approaches (e.g. adaptation neworks; Webster et al. 2017), preserve genetic diversity. Recruitment to the reef is another critical contributor towards coral fitness and population persistence (Ritson-Williams et al. 2009), and potential for high recruitment success is clearly an important factor for selecting an outplant reef. While the water quality parameters discussed above that promote adult colony success also likely contribute to juvenile health post-settlement, additional factors (e.g. settlement habitat, abundance of herbivores, prevalence of coral disease) are critical for recruitment to the reef (Ritson-Williams et al. 2009) and should also be considered when selecting outplant sites. Although sites in greatest need of restoration may not be “high fitness” sites, using corals from favorable sites or nurseries may still benefit the recipient population at a ‘low fitness’ reef for two reasons: 1) corals from a ‘high-fitness’ reef had higher reproductive success than native corals, likely boosting the fitness of the recipient population, and 2) introduction of bleaching resistant individuals should improve the fitness of that population during increasingly frequent marine heatwaves (Frölicher et al. 2018). Future work is needed to determine if either of these coral traits (high fitness and bleaching resistance) persists through the next spawning season or heatwave, or the next generation. Promisingly, putting corals in a good site improved their reproductive success, which could be beneficial for practitioners seeking to increase the genetic diversity of a recipient population (or rescue rare genets from a damaged or dying site), as transplanting these corals could be beneficial for the individual being moved without harming the recipient population. Thus, assisted gene flow could be another strategy for restoring and maintaining genetic diversity that also increases heat resistance of a population.

## Acknowledgements

This work would not have been possible without the support of volunteers in the field, and we would particularly like to thank Yanitza Grantcharska, Aileen Maldonado, Dyson Chee, and the HIMB boating and diving team led by Jason Jones. We also recognize technical help from Brian Glazer and Stanley Lio with sensor deployment.

