## Supplemental Materials for "Bleaching resistant corals retain heat tolerance following acclimatization to environmentally distinct reefs"

### Supplementary Methods

*Buoyant weight*: The density of the seawater was measured each day buoyant weight was measured using a glass standard of known density. For accurate conversion to dry weight, the skeletal density of each species was empirically determined from the buoyant weight and dry weight of tissue-free skeletons of each species according to the equations in (Davies, 1989). Mean skeletal density of *M. capitata* was 2.08 +/- 0.16 g $cm^{-3}$ (n=35) and *P. compressa* was 1.82 +/- 0.37 g $cm^{-3}$ (n=27). Calcification rates were normalized to the initial weight of the fragment resulting in units of % $day^{-1}$.

*Water flow:* Clod cards were molded in ice cube trays (4 x 3 x 3.5 cm) using Plaster of Paris mixed with water (2:1 plaster to water) and dried at 60°C for 3-5 days and weighed. Six clod cards were deployed at each reef site secured using embedded nylon bolts to transplant racks next to experimental corals and retrieved after 24-28 hours. Clod cards were then dried at 60°C for 3-5 days and weighed. The percent mass dissolution per hour at each reef site was calculated as a proxy for water flow rate.

*Sedimentation rate:* Sediment traps with a 7:1 ratio of tube height to mouth width (5 cm) diameter PVC tubes 42 cm in length (Storlazzi *et al.*, 2011). Sediment collected from traps was dried overnight at 60°C and weighed. Grams of sediment deposited per day at each reef site was calculated by dividing the dry weight of the sediment by the number of days the trap was deployed.

### Supplementary Figures

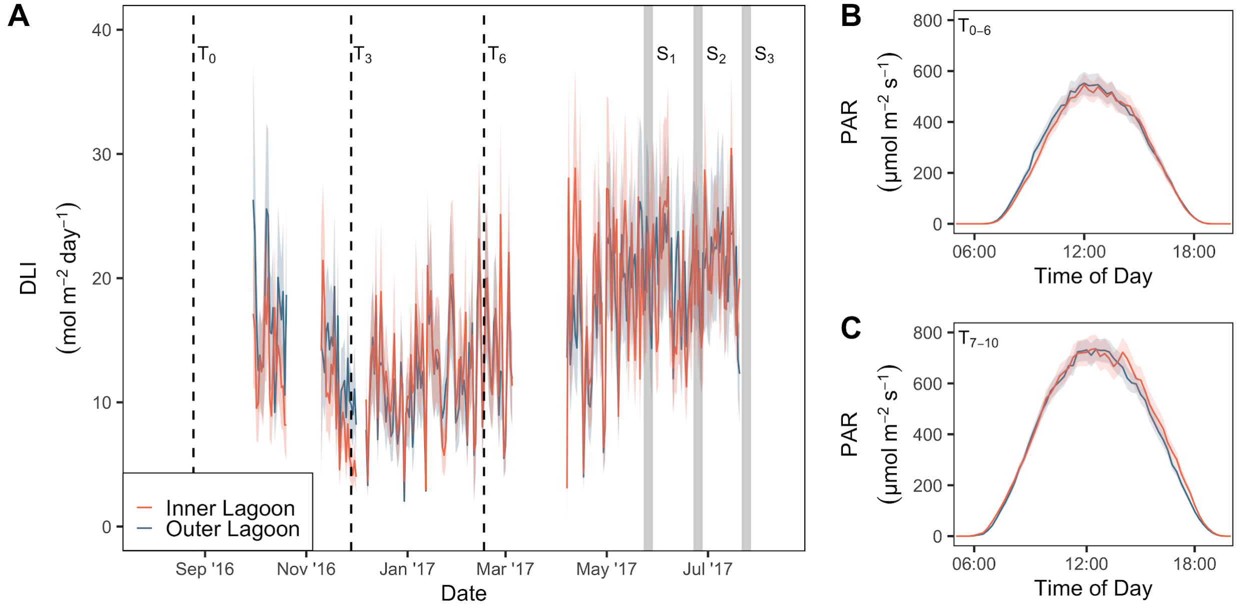


Figure S1. Light intensity of transplanted corals at the Inner and Outer Lagoon reefs (2 m depth). A) Mean daily light integral (DLI); B) Mean daily photosynthetically active radiation (PAR) for the first 6 months of acclimatization; C) Mean daily PAR for months 7-11 of reciprocal transplantation. Shaded regions in all graphs indicate 95% confidence intervals (N=3). Vertical dashed lines indicate the initiation of the transplant ($T_0$) followed by sampling time points after 3 months ($T_3$) and 6 months ($T_6$) of acclimatization. $S_{1-3}$ lines indicate coral spawning events where reproductive output was monitored.

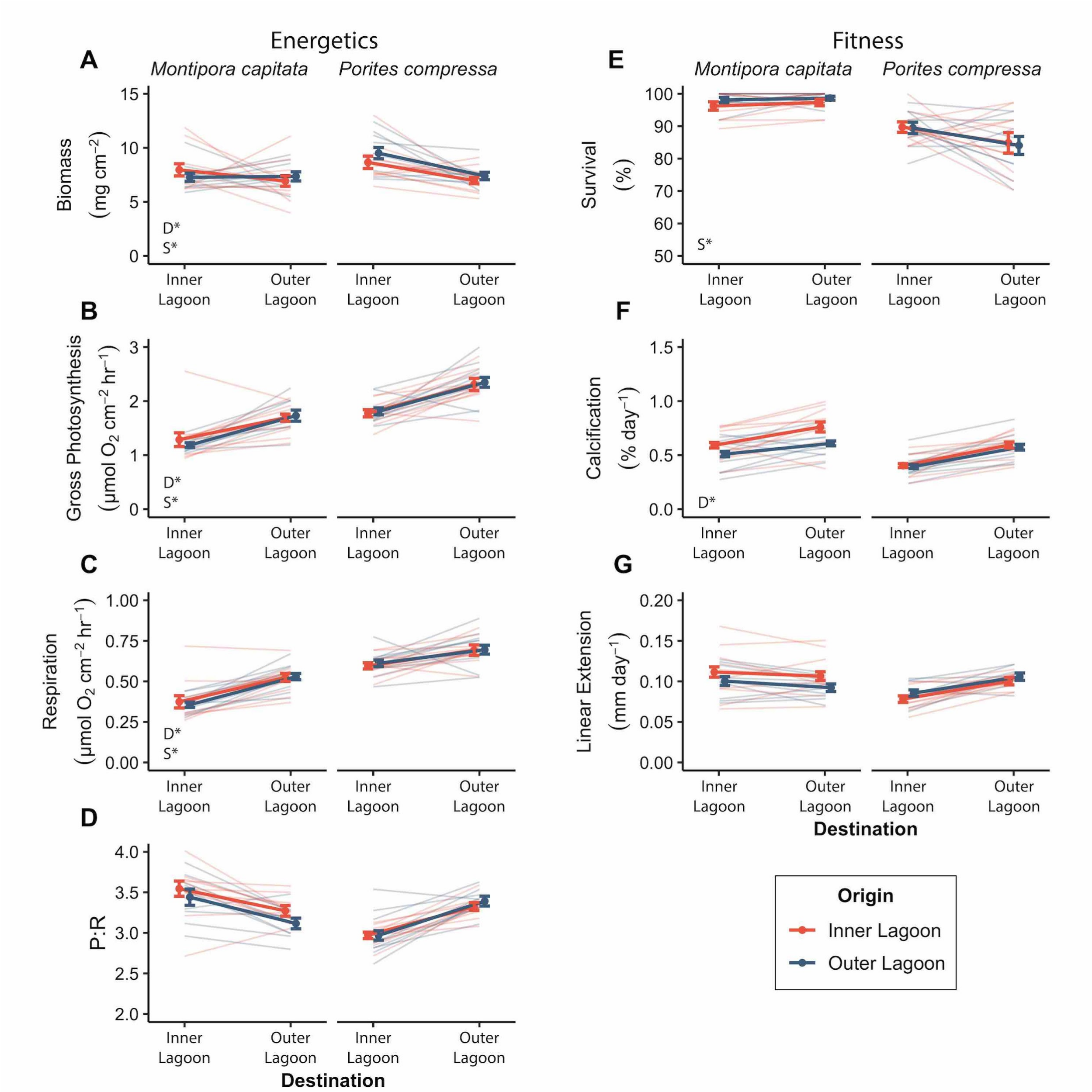


Figure S2. The effects of origin site and destination site on coral energetics (left) and fitness (right) after 3 months of reciprocal transplantation for *Montipora capitata* and *Porites compressa* between an Inner and Outer Lagoon reefs. A) Biomass; B) Gross photosynthesis; C) Respiration; D) Photosynthesis to respiration ratio (P:R); E) Survival; F) Calcification; and G) Linear extension. Bold lines indicate the mean of all genets (N=10) ± standard error of the mean (SEM); thin lines indicate mean response of each genet (N=5 ramets/genet).

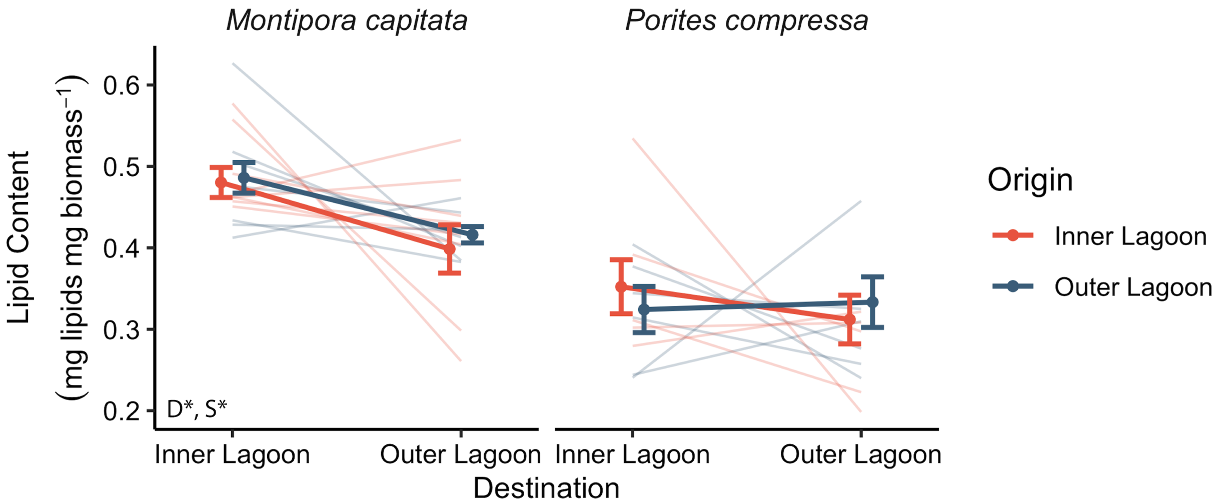


Figure S3. Coral tissue lipid content following six months of acclimatization for *M. capitata* (left) and *P. compressa* (right). Bold lines indicate the mean of all genets (N=10) ± standard error of the mean (SEM); thin lines indicate mean response of each genet (N=5 ramets/genet).

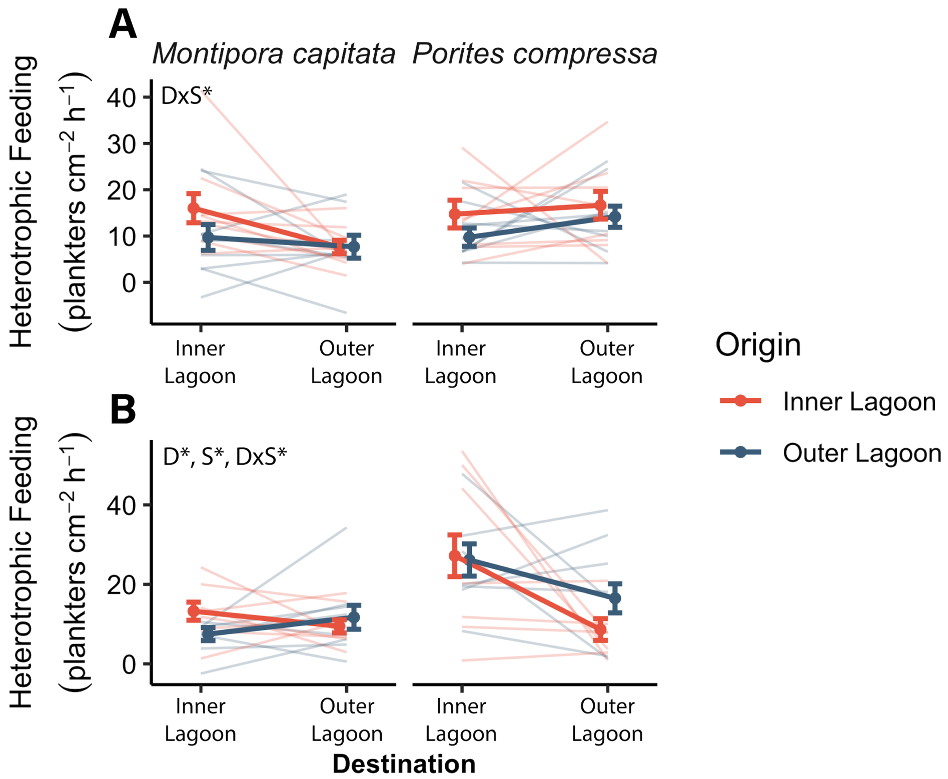


Figure S4. Heterotrophic feeding rates as measured by prey depletion of *M. capitata* (left) and *P. compressa* (right) after A) three months and B) six months of acclimatization. Bold lines indicate the mean of all genets (N=10) ± standard error of the mean (SEM); thin lines indicate mean response of each genet (N=5 fragments/genet).

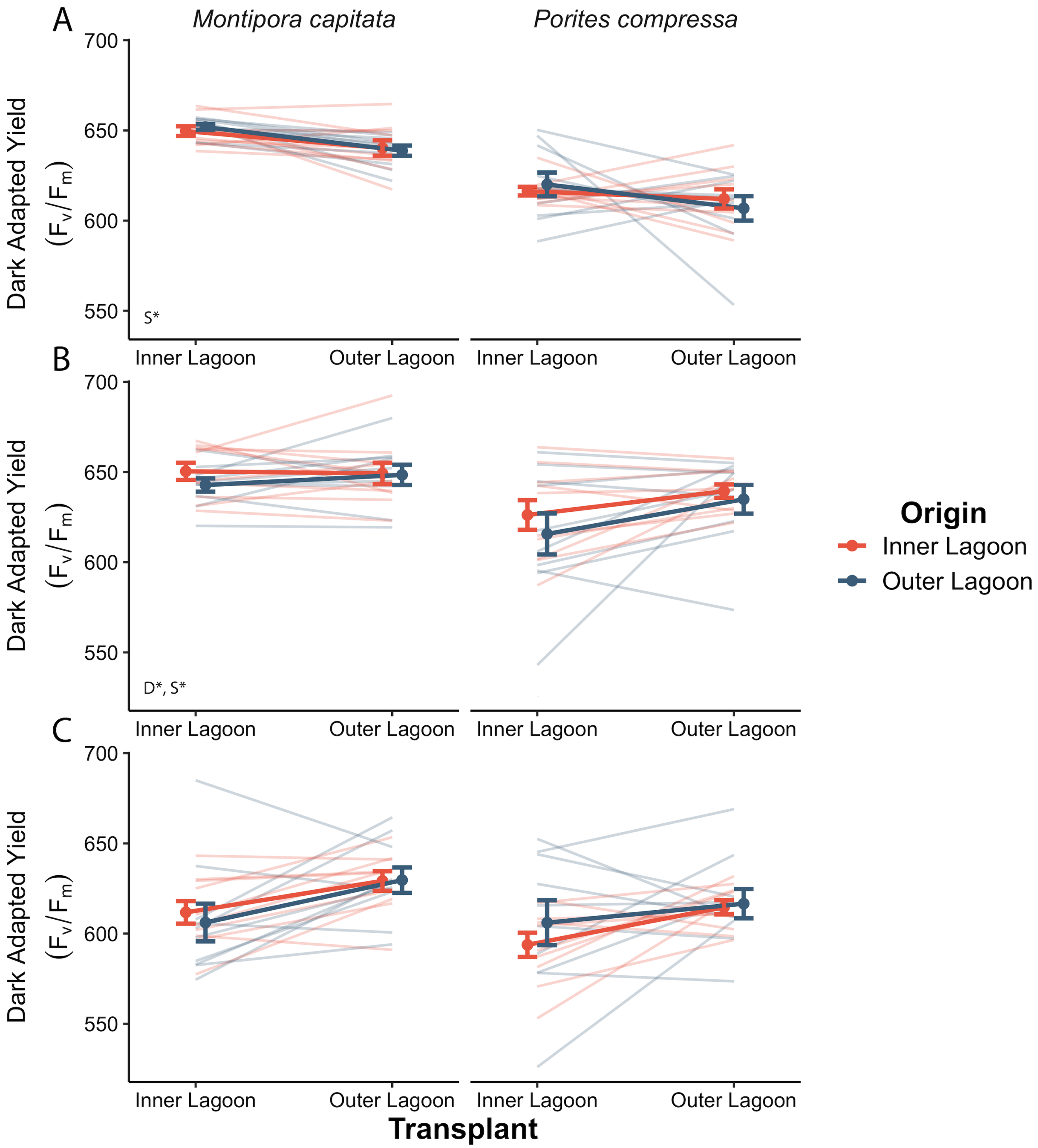


Figure S5. Photosynthetic efficiency of *M. capitata* (left) and *P. compressa* (right) as measured by dark adapted yield (Fv/Fm) following A) 1.5 months, B) 3 months, and C) 4.5 months of reciprocal transplantation. Thin lines indicate mean genet responses, thick lines indicate mean response of all genets for a given transplant history. Error bars indicate standard error.

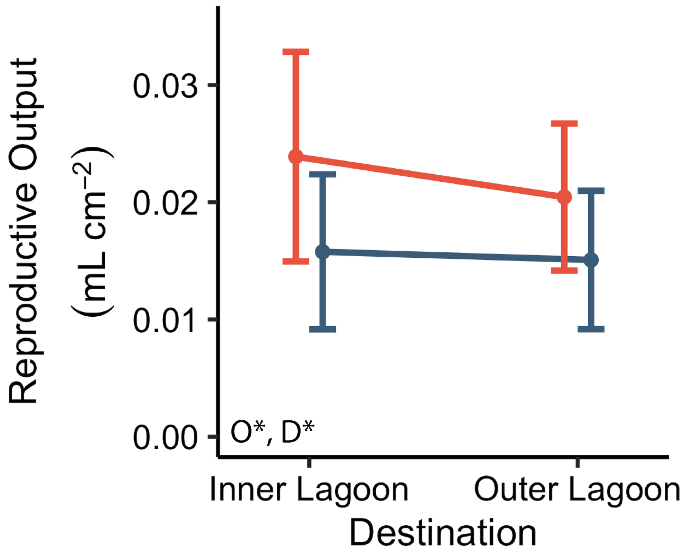


Figure S6. Fecundity of *M. capitata* as measured by reproductive output (volume of bundles released across the entire spawning season) following 9-10 months of reciprocal transplantation. N=6-10 genets; error bars indicate SEM.

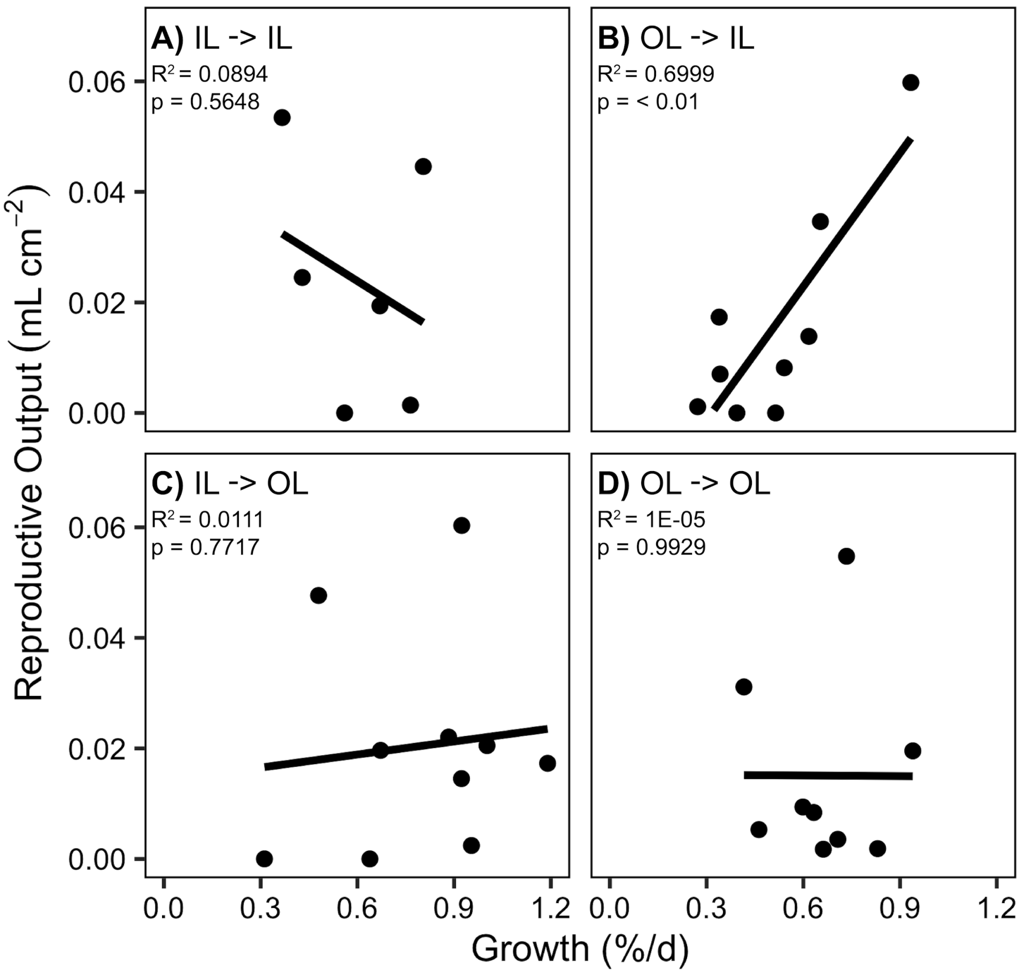


Figure S7. Relationship between growth rate and reproductive output for *M. capitata* for each transplant treatment: A) Inner Lagoon to Inner Lagoon, B) Outer Lagoon to Inner Lagoon, C) Inner Lagoon to Outer Lagoon, and D) Outer Lagoon to Outer Lagoon. N=6-10 genets per treatment; $R^2$ and p-values from linear regression (line) shown on each panel.

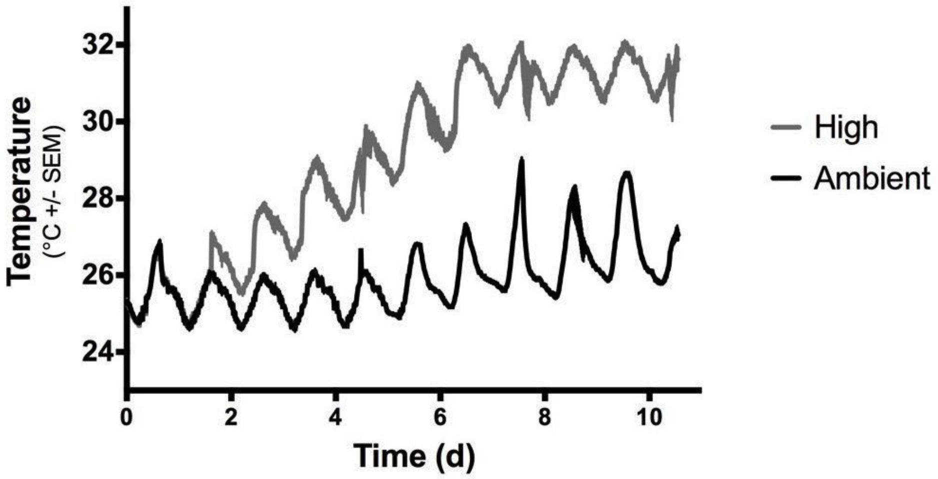


Figure S8. Mean temperature in ambient and high temperature tanks over the course of the acute heat stress experiment. The maximum daily temperature (32°C) is +4°C above the local MMM. N=4; error bars (SEM) lie within the width of the line.

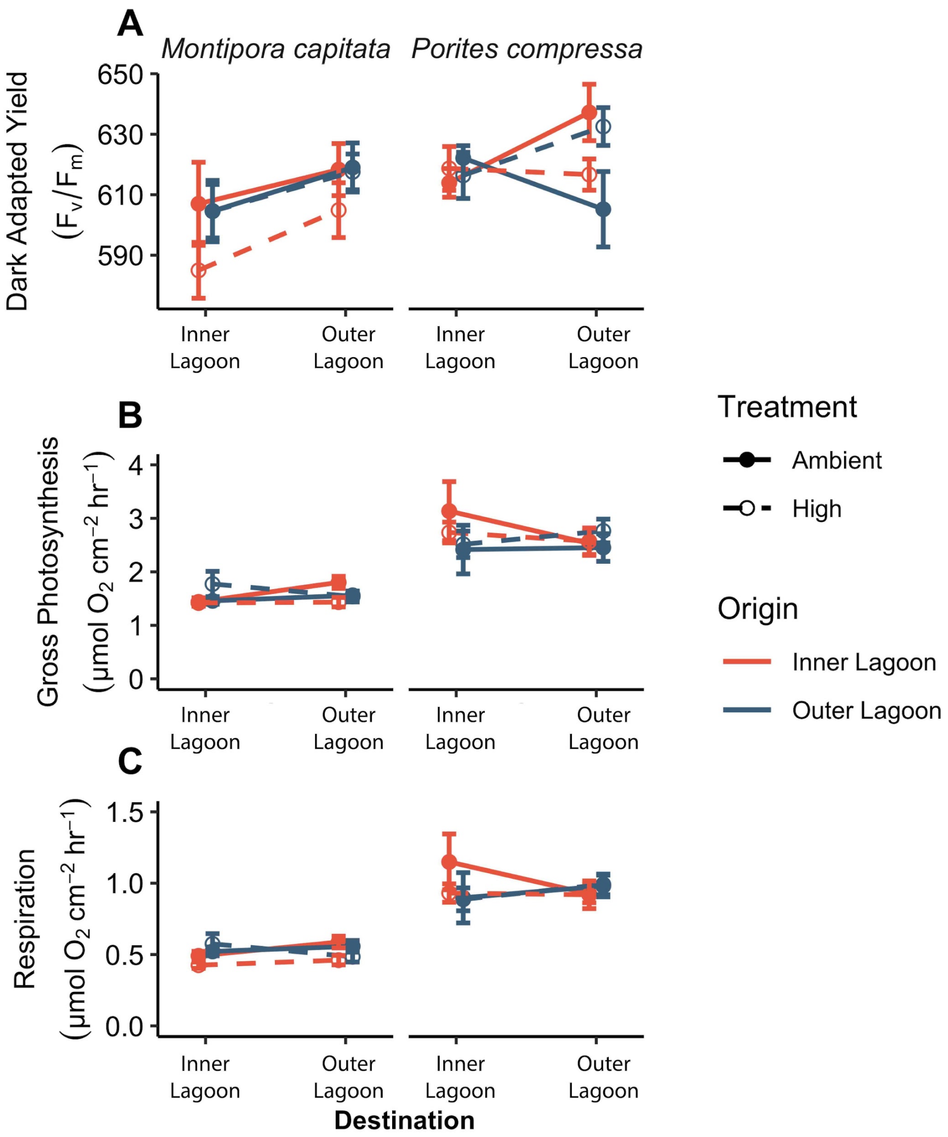


Figure S9. Coral physiology on Day 0 of the acute heat stress experiment. A) Dark adapted yield (Fv/Fm), B) gross photosynthesis rate, and C) dark respiration rate. N=10 genets per history per treatment; error bars indicate SEM.

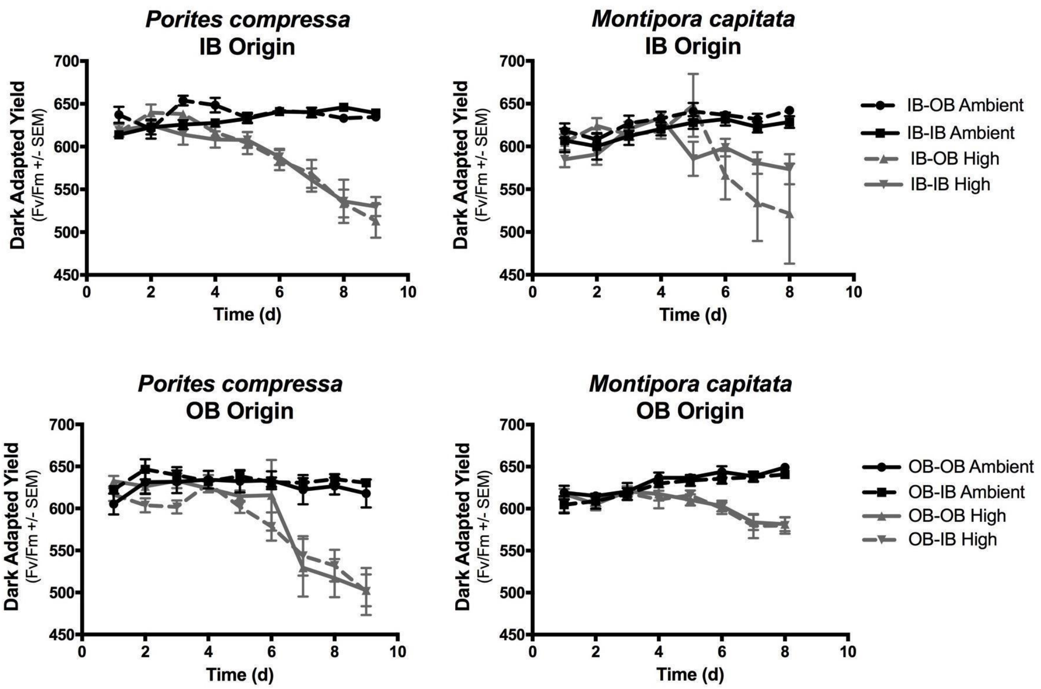


Figure S10. Time series of coral photosynthetic efficiency (dark adapted yield; Fv/Fm) across the acute temperature stress experiment. Error bars indicate SEM. Legend indicates origin-destination; IB, inner lagoon; OB, outer lagoon.

### Supplementary Tables

Table S1. Linear mixed effect model statistical analyses of coral response variables. Subset analyses indicate the factor for which separate analyses were conducted.

| **Response Variable** | **Subset Analysis** | **Model Class** | **Fixed Effects** | **Random Effects** |
|---|---|---|---|---|
| Linear Extension | Time point | lmer [1] | Origin Site; Destination Site; Species | Genotype; Rack |
| Calcification (square root transformed) | Time point | lmer [1] | Origin Site; Destination Site; Species | Genotype; Rack |
| Fitness Score | Species | lmer [1] | Origin Site; Destination Site | Genotype |
| Dark Adapted Yield (log transformed) | Time point | lmer [1] | Origin Site; Destination Site; Species | Genotype; Rack |
| Respiration | Time point | lmer [1] | Origin Site; Destination Site; Species | Genotype; Rack |
| Gross Photosynthesis | Time point | lmer [1] | Origin Site; Destination Site; Species | Genotype; Rack |
| Net Photosynthesis | Time point | lmer [1] | Origin Site; Destination Site; Species | Genotype; Rack |
| Heterotrophic Feeding (square root transformed) | Time point | lmer [1] | Origin Site; Destination Site; Species | Genotype |
| Biomass (log transformed) | Time point | lmer [1] | Origin Site; Destination Site; Species | Genotype; Rack |
| Lipid Content | 6-month time point | glmmTMB, beta family [2] | Origin Site; Destination Site; Species | Genotype; Rack |
| Spawning Activity | *M. capitata* only | glmer, binomial family [1] | Origin Site; Destination Site | Genotype; Date |
| Survivorship | Time point | betareg [3] | Origin Site; Destination Site; Species | |

1. *lme4* package (Bates et al. 2015); 2. *glmmTMB* package (Brooks et al. 2017); 3. *betareg* package (Cribari-Neto and Zeileis 2010); Significance of effects on lipid content, survivorship, and spawning activity determined by Chi Squared Type III ANOVA tests. All other responses analyzed by Type III Satterthwaite ANOVA tests.

Table S2. Statistical analysis of linear mixed effect models of coral biomass.

| Response Variable | Subset Analysis | Fixed Effect | SS | df | F | *p* |
|---|---|---|---|---|---|---|
| Biomass | 3-month | Origin | 0.38 | 1, 355 | 2.20 | 0.14 |
| | | **Destination** | **1.57** | **1, 372** | **9.15** | **<0.01** |
| | | **Species** | **1.41** | **1, 355** | **8.24** | **<0.01** |
| | | Origin x Destination | 0.00 | 1, 372 | 0.00 | 0.99 |
| | | Origin x Species | 0.08 | 1, 355 | 0.05 | 0.83 |
| | | Destination x Species | 0.62 | 1, 372 | 3.62 | 0.06 |
| | | Origin x Destination x Species | 0.08 | 1, 372 | 0.49 | 0.48 |
| | 6-month | Origin | 0.32 | 1, 350 | 2.02 | 0.16 |
| | | Destination | 0.01 | 1, 350 | 0.07 | 0.78 |
| | | **Species** | **6.26** | **1, 350** | **39.91** | **<0.01** |
| | | Origin x Destination | 0.08 | 1, 350 | 0.57 | 0.45 |
| | | Origin x Species | 0.00 | 1, 350 | 0.00 | 0.97 |
| | | **Destination x Species** | **2.13** | **1, 350** | **13.56** | **<0.01** |
| | | Origin x Destination x Species | 0.01 | 1, 350 | 0.06 | 0.81 |

SS = Sum of Squares; df = degrees of freedom (numerator, denominator); F = F statistic; bold indicates statistical significance (p<0.05).

Table S3. Summary statistics of coral biomass.

| **Species** | **Origin** | **Destination** | **Biomass (mg cm$^{-2}$)** | | | | | |
|---|---|---|---|---|---|---|---|---|
| | | | 3 months | | | 6 months | | |
| | | | N | mean | sem | N | mean | sem |
| *Montipora capitata* | Inner Lagoon | Inner Lagoon | 48 | 7.96 | 0.56 | 46 | 5.65 | 0.49 |
| | | Outer Lagoon | 48 | 6.93 | 0.49 | 40 | 4.51 | 0.25 |
| | Outer Lagoon | Inner Lagoon | 48 | 7.28 | 0.37 | 51 | 5.63 | 0.28 |
| | | Outer Lagoon | 49 | 7.35 | 0.42 | 50 | 4.99 | 0.32 |
| *Porites Compressa* | Inner Lagoon | Inner Lagoon | 48 | 8.66 | 0.58 | 43 | 6.28 | 0.45 |
| | | Outer Lagoon | 46 | 6.96 | 0.28 | 46 | 7.01 | 0.36 |
| | Outer Lagoon | Inner Lagoon | 49 | 9.51 | 0.52 | 40 | 6.36 | 0.39 |
| | | Outer Lagoon | 44 | 7.39 | 0.25 | 42 | 7.62 | 0.37 |

Table S4. Statistical analysis of linear mixed effect models of coral lipid content.

| Response Variable | Subset Analysis | Fixed Effect | df | Chi Squared | *p* |
|---|---|---|---|---|---|
| Lipid Content | 6-month | Origin | 1 | 0.03 | 0.85 |
| | | **Destination** | **1** | **6.09** | **0.01** |
| | | **Species** | **1** | **12.75** | **<0.01** |
| | | Origin x Destination | 1 | 0.75 | 0.75 |
| | | Origin x Species | 1 | 0.45 | 0.45 |
| | | Destination x Species | 1 | 0.47 | 0.47 |
| | | Origin x Destination x Species | 1 | 0.56 | 0.56 |

df = degrees of freedom; bold indicates statistical significance (p<0.05).

Table S5. Summary statistics of coral lipid content.

| **Species** | **Origin** | **Destination** | **Lipid Content (mg lipids mg biomass$^{-1}$)** 6 months N | mean | sem |
|---|---|---|---|---|---|
| *Montipora capitata* | Inner Lagoon | Inner Lagoon | 9 | 0.48 | 0.02 |
| | | Outer Lagoon | 10 | 0.40 | 0.03 |
| | Outer Lagoon | Inner Lagoon | 11 | 0.49 | 0.02 |
| | | Outer Lagoon | 10 | 0.42 | 0.01 |
| *Porites Compressa* | Inner Lagoon | Inner Lagoon | 11 | 0.35 | 0.03 |
| | | Outer Lagoon | 9 | 0.31 | 0.03 |
| | Outer Lagoon | Inner Lagoon | 8 | 0.32 | 0.03 |
| | | Outer Lagoon | 11 | 0.33 | 0.03 |

Table S6. Statistical analysis of linear mixed effect models of coral heterotrophic feeding.

| **Response Variable** | **Subset Analysis** | **Fixed Effect** | **SS** | **df** | **F** | ***p*** |
|---|---|---|---|---|---|---|
| Heterotrophic Feeding | 3-month | Origin | 2.89 | 1, 32 | 3.76 | 0.06 |
| | | Destination | 0.35 | 1, 34 | 0.46 | 0.50 |
| | | Species | 2.87 | 1, 32 | 3.73 | 0.06 |
| | | Origin x Destination | 1.06 | 1, 34 | 1.37 | 0.25 |
| | | Origin x Species | 0.00 | 1, 32 | 0.00 | 0.96 |
| | | **Destination x Species** | **4.35** | **1, 34** | **5.66** | **0.02** |
| | | Origin x Destination x Species | 0.10 | 1, 34 | 0.13 | 0.72 |
| | 6-month | Origin | 0.10 | 1, 37 | 0.10 | 0.75 |
| | | **Destination** | **8.28** | **1, 52** | **8.20** | **<0.01** |
| | | **Species** | **10.92** | **1, 37** | **10.81** | **<0.01** |
| | | Origin x Destination | 3.52 | 1, 52 | 3.48 | 0.06 |
| | | Origin x Species | 1.64 | 1, 37 | 1.63 | 0.21 |
| | | **Destination x Species** | **9.26** | **1, 52** | **9.18** | **<0.01** |
| | | Origin x Destination x Species | 0.00 | 1, 52 | 0.00 | 0.97 |

SS = Sum of Squares; df = degrees of freedom (numerator, denominator); F = F statistic; bold indicates statistical significance (p<0.05).

Table S7. Summary statistics of coral heterotrophic feeding.

| **Species** | **Origin** | **Destination** | **Heterotrophic Feeding (prey consumed $cm^{-2}$ $h^{-1}$)** | | | | | |
|---|---|---|---|---|---|---|---|---|
| | | | 3 months | | | 6 months | | |
| | | | N | mean | sem | N | mean | sem |
| *Montipora capitata* | Inner Lagoon | Inner Lagoon | 10 | 16.01 | 3.15 | 9 | 13.25 | 2.28 |
| | | Outer Lagoon | 9 | 7.63 | 1.45 | 10 | 9.45 | 1.65 |

| | | | | | | | | |
|---|---|---|---|---|---|---|---|---|
| | Outer Lagoon | Inner Lagoon | 10 | 9.69 | 2.79 | 11 | 7.50 | 1.65 |
| | | Outer Lagoon | 9 | 7.69 | 2.48 | 10 | 11.71 | 3.01 |
| *Porites Compressa* | Inner Lagoon | Inner Lagoon | 8 | 14.73 | 2.99 | 11 | 27.20 | 5.25 |
| | | Outer Lagoon | 10 | 16.65 | 2.98 | 9 | 8.64 | 2.73 |
| | Outer Lagoon | Inner Lagoon | 11 | 9.74 | 1.99 | 9 | 26.14 | 4.05 |
| | | Outer Lagoon | 10 | 14.16 | 2.29 | 11 | 16.49 | 3.66 |

Table S8. Statistical analysis of linear mixed effect models of coral dark-adapted yield. Dark-adapted yield data were log transformed to meet assumptions of normality.

| **Response Variable** | **Subset Analysis** | **Fixed Effect** | **SS** | **df** | **F** | ***p*** |
|---|---|---|---|---|---|---|
| Dark Adapted Yield | 1.5-month | Origin | 0.00 | 1, 762 | 0.00 | 0.93 |
| | | Destination | 0.02 | 1, 5 | 4.43 | 0.09 |
| | | **Species** | **0.13** | **1, 5** | **30.52** | **<0.01** |
| | | Origin x Destination | 0.00 | 1, 762 | 0.24 | 0.62 |
| | | Origin x Species | 0.00 | 1, 762 | 0.44 | 0.51 |
| | | Destination x Species | 0.00 | 1, 5 | 0.02 | 0.89 |
| | | Origin x Destination x Species | 0.00 | 1, 762 | 0.32 | 0.57 |
| | 3-month | Origin | 0.00 | 1, 35 | 0.44 | 0.51 |
| | | **Destination** | **0.02** | **1, 302** | **5.94** | **0.02** |
| | | **Species** | **0.04** | **1, 35** | **9.31** | **<0.01** |
| | | Origin x Destination | 0.00 | 1, 336 | 0.45 | 0.50 |
| | | Origin x Species | 0.00 | 1, 35 | 0.00 | 0.98 |
| | | Destination x Species | 0.01 | 1, 302 | 3.47 | 0.06 |
| | | Origin x Destination x Species | 0.00 | 1, 336 | 0.04 | 0.85 |
| | 4.5-month | Origin | 0.00 | 1, 35 | 0.34 | 0.57 |
| | | Destination | 0.00 | 1, 0.5 | 0.09 | 0.84 |
| | | Species | 0.00 | 1, 0.4 | 0.12 | 0.84 |
| | | Origin x Destination | 0.00 | 1, 332 | 0.31 | 0.58 |
| | | Origin x Species | 0.00 | 1, 35 | 0.89 | 0.35 |
| | | Destination x Species | 0.00 | 1, 0.5 | 0.24 | 0.76 |
| | | Origin x Destination x Species | 0.00 | 1, 332 | 0.71 | 0.40 |

SS = Sum of Squares; df = degrees of freedom (numerator, denominator); F = F statistic; bold indicates statistical significance ($p<0.05$). Models violated assumption of homogeneity of variance.

Table S9. Summary statistics of coral dark adapted yield.

| Species | Origin | Destination | Dark Adapted Yield | | | | | | | | |
|---|---|---|---|---|---|---|---|---|---|---|---|
| | | | 1.5 months | | | 3 months | | | 4.5 months | | |
| | | | N | mean | sem | N | mean | sem | N | mean | sem |
| *Montipora capitata* | Inner Lagoon | Inner Lagoon | 99 | 649.58 | 2.68 | 51 | 648.88 | 3.87 | 50 | 610.80 | 8.00 |
| | | Outer Lagoon | 98 | 636.78 | 3.52 | 46 | 646.98 | 3.60 | 46 | 631.52 | 5.38 |
| | Outer Lagoon | Inner Lagoon | 101 | 651.64 | 2.73 | 52 | 641.88 | 5.27 | 52 | 606.83 | 6.61 |
| | | Outer Lagoon | 101 | 638.52 | 2.26 | 50 | 647.80 | 5.41 | 49 | 629.86 | 5.35 |
| *Porites Compress a* | Inner Lagoon | Inner Lagoon | 96 | 616.08 | 3.28 | 44 | 625.48 | 8.31 | 45 | 594.27 | 5.59 |
| | | Outer Lagoon | 93 | 608.35 | 5.23 | 44 | 640.18 | 4.78 | 44 | 615.90 | 4.15 |
| | Outer Lagoon | Inner Lagoon | 96 | 619.04 | 4.96 | 40 | 619.38 | 8.33 | 40 | 609.73 | 6.82 |

| | | | | | | | | | | | |
|---|---|---|---|---|---|---|---|---|---|---|---|
| | | Outer Lagoon | 90 | 604.06 | 5.71 | 42 | 639.90 | 6.33 | 42 | 620.00 | 5.57 |

Table S10. Statistical analysis of linear mixed effect models of coral respiration and photosynthesis rates.

| **Response Variable** | **Subset Analysis** | **Fixed Effect** | **SS** | **df** | **F** | ***p*** |
|---|---|---|---|---|---|---|
| Respiration | 3-month | Origin | 2.0e-08 | 1, 35 | 0.00 | 0.96 |
| | | **Destination** | **3.8e-04** | **1, 311** | **52.42** | **<0.01** |
| | | **Species** | **5.2e-04** | **1, 35** | **74.03** | **<0.01** |
| | | Origin x Destination | 6.1e-07 | 1, 312 | 0.09 | 0.77 |
| | | Origin x Species | 6.5e-07 | 1, 35 | 0.09 | 0.77 |
| | | Destination x Species | 2.7e-05 | 1, 311 | 3.81 | 0.05 |
| | | Origin x Destination x Species | 1.6e-06 | 1, 312 | 0.23 | 0.63 |
| | 6-month | Origin | 4.5e-07 | 1, 35 | 0.06 | 0.80 |
| | | Destination | 9.0e-09 | 1, 0.7 | 0.00 | 0.98 |
| | | Species | 2.0e-05 | 1, 0.5 | 2.81 | 0.48 |
| | | **Origin x Destination** | **4.5e-05** | **1, 317** | **6.36** | **0.01** |
| | | Origin x Species | 0.00 | 1, 35 | 0.00 | 0.99 |
| | | Destination x Species | 2.4e-06 | 1, 0.7 | 0.34 | 0.69 |
| | | Origin x Destination x Species | 9.6e-06 | 1, 317 | 1.36 | 0.25 |
| Gross Photosynthesis | 3-month | Origin | 2.0e-07 | 1, 35 | 0.00 | 0.97 |
| | | **Destination** | **6.1e-03** | **1, 311** | **67.74** | **<0.01** |
| | | **Species** | **5.5e-03** | **1, 35** | **60.87** | **<0.01** |
| | | Origin x Destination | 4.4e-05 | 1, 311 | 0.50 | 0.48 |
| | | Origin x Species | 1.7e-05 | 1, 35 | 0.19 | 0.67 |
| | | Destination x Species | 2.8e-05 | 1, 311 | 0.31 | 0.58 |
| | | Origin x Destination x Species | 2.1e-05 | 1, 311 | 0.24 | 0.62 |
| | 6-month | Origin | 2.2e-07 | 1, 34 | 0.00 | 0.96 |
| | | Destination | 5.0e-06 | 1, 0.7 | 0.06 | 0.86 |
| | | Species | 1.3e-04 | 1, 0.48 | 1.65 | 0.55 |
| | | **Origin x Destination** | **4.7e-04** | **1, 319** | **5.97** | **0.02** |
| | | Origin x Species | 1.2e-05 | 1, 34 | 0.16 | 0.69 |
| | | Destination x Species | 5.8e-05 | 1, 0.7 | 0.70 | 0.60 |
| | | Origin x Destination x Species | 6.6e-05 | 1, 319 | 0.83 | 0.36 |
| P:R Ratio | 3-month | Origin | 0.02 | 1, 39 | 0.04 | 0.85 |
| | | Destination | 1.11 | 1, 1.5 | 1.83 | 0.34 |
| | | Species | 4.41 | 1, 2 | 7.23 | 0.11 |
| | | Origin x Destination | 0.43 | 1, 318 | 0.71 | 0.40 |
| | | Origin x Species | 0.02 | 1, 39 | 0.03 | 0.86 |
| | | Destination x Species | 6.67 | 1, 2 | 10.98 | 0.11 |
| | | Origin x Destination x Species | 0.11 | 1, 318 | 0.18 | 0.67 |
| | 6-month | Origin | 0.25 | 1, 36 | 1.71 | 0.19 |
| | | **Destination** | **2.19** | **1, 317** | **15.05** | **<0.01** |
| | | **Species** | **7.03** | **1, 36** | **48.23** | **<0.01** |
| | | Origin x Destination | 0.03 | 1, 317 | 0.19 | 0.66 |
| | | Origin x Species | 0.28 | 1, 36 | 1.91 | 0.18 |
| | | **Destination x Species** | **2.42** | **1, 317** | **16.64** | **<0.01** |
| | | Origin x Destination x Species | 0.04 | 1, 317 | 0.27 | 0.60 |

SS = Sum of Squares; df = degrees of freedom (numerator, denominator); F = F statistic; bold indicates statistical significance ($p<0.05$).

Table S11. Summary statistics of coral gross photosynthesis.

| Species | Origin | Destination | Gross Photosynthesis (umol cm$^{-2}$ min$^{-1}$) | | | | | |
|---|---|---|---|---|---|---|---|---|
| | | | 3 months | | | 6 months | | |
| | | | N | mean | sem | N | mean | sem |
| *Montipora capitata* | Inner Lagoon | Inner Lagoon | 44 | 0.021 | 0.002 | 43 | 0.019 | 0.002 |
| | | Outer Lagoon | 42 | 0.028 | 0.001 | 40 | 0.020 | 0.001 |
| | Outer Lagoon | Inner Lagoon | 45 | 0.020 | 0.001 | 38 | 0.015 | 0.001 |
| | | Outer Lagoon | 45 | 0.029 | 0.002 | 45 | 0.022 | 0.001 |
| *Porites Compressa* | Inner Lagoon | Inner Lagoon | 42 | 0.030 | 0.001 | 43 | 0.023 | 0.001 |
| | | Outer Lagoon | 44 | 0.038 | 0.002 | 46 | 0.030 | 0.001 |
| | Outer Lagoon | Inner Lagoon | 45 | 0.030 | 0.001 | 39 | 0.025 | 0.001 |
| | | Outer Lagoon | 42 | 0.039 | 0.002 | 42 | 0.032 | 0.002 |

Table S12. Summary statistics of coral respiration.

| Species | Origin | Destination | Respiration (umol cm$^{-2}$ min$^{-1}$) | | | | | |
|---|---|---|---|---|---|---|---|---|
| | | | 3 months | | | 6 months | | |
| | | | N | mean | sem | N | mean | sem |
| *Montipora capitata* | Inner Lagoon | Inner Lagoon | 44 | 0.006 | 0.001 | 43 | 0.006 | 0.001 |
| | | Outer Lagoon | 42 | 0.009 | 0.000 | 40 | 0.006 | 0.000 |
| | | Inner Lagoon | 45 | 0.006 | 0.000 | 48 | 0.005 | 0.000 |

| | | | | | | | | |
|---|---|---|---|---|---|---|---|---|
| | Outer Lagoon | Outer Lagoon | 45 | 0.009 | 0.000 | 45 | 0.007 | 0.000 |
| *Porites Compressa* | Inner Lagoon | Inner Lagoon | 42 | 0.010 | 0.000 | 43 | 0.009 | 0.000 |
| | | Outer Lagoon | 44 | 0.012 | 0.001 | 46 | 0.010 | 0.000 |
| | Outer Lagoon | Inner Lagoon | 45 | 0.010 | 0.000 | 39 | 0.009 | 0.000 |
| | | Outer Lagoon | 42 | 0.012 | 0.000 | 42 | 0.010 | 0.000 |

Table S13. Summary statistics of coral P:R.

| **Species** | **Origin** | **Destination** | **Photosynthesis : Respiration** | | | | | |
|---|---|---|---|---|---|---|---|---|
| | | | 3 months | | | 6 months | | |
| | | | N | mean | sem | N | mean | sem |
| *Montipora capitata* | Inner Lagoon | Inner Lagoon | 44 | 3.55 | 0.09 | 43 | 3.44 | 0.08 |
| | | Outer Lagoon | 42 | 3.27 | 0.07 | 40 | 3.44 | 0.07 |
| | Outer Lagoon | Inner Lagoon | 45 | 3.44 | 0.10 | 48 | 3.29 | 0.07 |
| | | Outer Lagoon | 44 | 3.12 | 0.07 | 45 | 3.28 | 0.06 |
| *Porites Compressa* | Inner Lagoon | Inner Lagoon | 42 | 2.97 | 0.04 | 43 | 2.84 | 0.04 |
| | | Outer Lagoon | 44 | 3.33 | 0.05 | 46 | 3.13 | 0.06 |
| | Outer Lagoon | Inner Lagoon | 45 | 2.97 | 0.06 | 39 | 2.81 | 0.05 |
| | | Outer Lagoon | 42 | 3.39 | 0.06 | 42 | 3.17 | 0.05 |

Table S14. Statistical analysis of linear mixed effect models of coral survival and reproduction.

| Response Variable | Subset Analysis | Fixed Effect | df | Chi Squared | *p* |
|---|---|---|---|---|---|
| Spawning Activity | *M. capitata* | **Origin** | **1** | **6.31** | **0.01** |
| | | **Destination** | **1** | **4.52** | **0.03** |
| | | Origin x Destination | 1 | 3.35 | 0.07 |
| Survivorship | 3-months | Origin | 1 | 2.70 | 0.10 |
| | | Destination | 1 | 0.10 | 0.75 |
| | | **Species** | **1** | **4.85** | **0.03** |
| | | Origin x Destination | 1 | 2.16 | 0.14 |
| | | Origin x Species | 1 | 0.26 | 0.61 |
| | | Destination x Species | 1 | 2.55 | 0.11 |
| | | Origin x Destination x Species | 1 | 0.33 | 0.57 |
| | 6-months | Origin | 1 | 0.33 | 0.56 |
| | | Destination | 1 | 0.02 | 0.90 |
| | | **Species** | **1** | **9.58** | **<0.01** |
| | | Origin x Destination | 1 | 0.03 | 0.87 |
| | | Origin x Species | 1 | 1.78 | 0.18 |
| | | Destination x Species | 1 | 1.87 | 0.17 |
| | | Origin x Destination x Species | 1 | 0.11 | 0.74 |

df = degrees of freedom; bold indicates statistical significance (p<0.05).

Table S15. Summary statistics of coral survivorship.

| | | | **Survivorship** | | | | | |
|---|---|---|---|---|---|---|---|---|
| **Species** | **Origin** | **Destination** | 3 months | | | 6 months | | |
| | | | N | mean | sem | N | mean | sem |
| *Montipora capitata* | Inner Lagoon | Inner Lagoon | 10 | 0.87 | 0.05 | 10 | 0.96 | 0.01 |
| | | Outer Lagoon | 10 | 0.95 | 0.02 | 10 | 0.97 | 0.01 |
| | Outer Lagoon | Inner Lagoon | 10 | 0.91 | 0.03 | 10 | 0.98 | 0.01 |
| | | Outer Lagoon | 10 | 0.96 | 0.02 | 10 | 0.99 | 0.01 |
| *Porites Compressa* | Inner Lagoon | Inner Lagoon | 10 | 0.88 | 0.02 | 10 | 0.9 | 0.01 |
| | | Outer Lagoon | 10 | 0.81 | 0.04 | 10 | 0.85 | 0.03 |

| | | | | | | | |
|---|---|---|---|---|---|---|---|
| Outer Lagoon | Inner Lagoon | 10 | 0.88 | 0.03 | 10 | 0.89 | 0.02 |
| | Outer Lagoon | 10 | 0.81 | 0.04 | 10 | 0.84 | 0.03 |

*N indicates parental colony*

Table S16. Statistical analysis of linear mixed effect models of the coral growth metrics linear extension and calcification.

| Response Variable | Subset Analysis | Fixed Effect | SS | df | F | *p* |
|---|---|---|---|---|---|---|
| Linear Extension | 0-3-months | Origin | 0.0006 | 1, 36 | 0.11 | 0.75 |
| | | Destination | 0.0029 | 1, 6 | 0.53 | 0.49 |
| | | Species | 0.0018 | 1, 7 | 0.32 | 0.59 |
| | | Origin x Destination | 0.0018 | 1, 686 | 0.31 | 0.58 |
| | | Origin x Species | 0.0160 | 1, 36 | 2.87 | 0.10 |
| | | Destination x Species | 0.0060 | 1, 6 | 1.07 | 0.34 |
| | | Origin x Destination x Species | 0.0000 | 1, 686 | 0.01 | 0.94 |
| | 0-6-months | Origin | 0.000 | 1, 36 | 0.14 | 0.70 |
| | | Destination | 0.001 | 1, 0.1 | 1.05 | 0.83 |
| | | Species | 0.016 | 1, 0.1 | 15.55 | 0.71 |
| | | Origin x Destination | 0.000 | 1, 327 | 0.05 | 0.82 |
| | | Origin x Species | 0.001 | 1, 36 | 6.65 | 0.42 |
| | | Destination x Species | 0.009 | 1, 0.1 | 8.89 | 0.76 |
| | | Origin x Destination x Species | 0.000 | 1, 327 | 0.01 | 0.93 |
| Calcification | 0-3-months | Origin | 0.04 | 1, 36 | 1.83 | 0.18 |
| | | **Destination** | **0.25** | **1, 4** | **11.15** | **0.03** |
| | | Species | 0.09 | 1, 9 | 3.98 | 0.08 |
| | | Origin x Destination | 0.01 | 1, 704 | 0.44 | 0.51 |
| | | Origin x Species | 0.03 | 1, 36 | 1.19 | 0.28 |

| | | | | | | |
|---|---|---|---|---|---|---|
| | | Destination x Species | 0.01 | 1, 4 | 0.26 | 0.63 |
| | | Origin x Destination x Species | 0.00 | 1,704 | 0.15 | 0.70 |
| | 0-6-months | Origin | 0.05 | 1, 36 | 2.00 | 0.16 |
| | | Destination | 0.05 | 1, 314 | 1.79 | 0.18 |
| | | Species | 0.01 | 1, 340 | 0.43 | 0.51 |
| | | Origin x Destination | 0.01 | 1, 321 | 0.19 | 0.66 |
| | | Origin x Species | 0.01 | 1, 36 | 0.50 | 0.48 |
| | | Destination x Species | 0.01 | 1, 314 | 0.22 | 0.64 |
| | | Origin x Destination x Species | 0.01 | 1, 321 | 0.51 | 0.48 |

SS = Sum of Squares; df = degrees of freedom (numerator, denominator); F = F statistic; bold indicates statistical significance (p<0.05).

Table S17. Summary statistics of coral calcification (buoyant weight).

| Species | Origin | Destination | Buoyant Weight (% day$^{-1}$) 0-3 months N | | | 0-6 months N | | |
|---|---|---|---|---|---|---|---|---|
| | | | N | mean | sem | N | mean | sem |
| *Montipora capitata* | Inner Lagoon | Inner Lagoon | 97 | 0.59 | 0.03 | 46 | 0.59 | 0.04 |
| | | Outer Lagoon | 94 | 0.76 | 0.05 | 45 | 0.83 | 0.06 |
| | | Inner Lagoon | 100 | 0.51 | 0.02 | 49 | 0.51 | 0.04 |

| | | | | | | | | |
|---|---|---|---|---|---|---|---|---|
| | Outer Lagoon | Outer Lagoon | 99 | 0.61 | 0.02 | 49 | 0.67 | 0.04 |
| *Porites Compressa* | Inner Lagoon | Inner Lagoon | 92 | 0.40 | 0.02 | 43 | 0.36 | 0.03 |
| | | Outer Lagoon | 92 | 0.59 | 0.03 | 45 | 0.71 | 0.04 |
| | Outer Lagoon | Inner Lagoon | 89 | 0.40 | 0.02 | 40 | 0.33 | 0.03 |
| | | Outer Lagoon | 87 | 0.57 | 0.03 | 41 | 0.68 | 0.04 |

Table S18. Summary statistics of coral linear extension.

| **Species** | **Origin** | **Destination** | **Linear Extension (mm day$^{-1}$)** | | | | | |
|---|---|---|---|---|---|---|---|---|
| | | | 0-3 months | | | 0-6 months | | |
| | | | N | mean | sem | N | mean | sem |
| *Montipora capitata* | Inner Lagoon | Inner Lagoon | 90 | 0.11 | 0.006 | 48 | 0.12 | 0.006 |
| | | Outer Lagoon | 87 | 0.11 | 0.005 | 46 | 0.11 | 0.005 |
| | Outer Lagoon | Inner Lagoon | 97 | 0.10 | 0.005 | 50 | 0.12 | 0.005 |
| | | Outer Lagoon | 95 | 0.09 | 0.005 | 47 | 0.10 | 0.004 |
| *Porites Compressa* | Inner Lagoon | Inner Lagoon | 92 | 0.08 | 0.004 | 42 | 0.05 | 0.004 |
| | | Outer Lagoon | 92 | 0.10 | 0.005 | 45 | 0.09 | 0.005 |
| | Outer Lagoon | Inner Lagoon | 87 | 0.09 | 0.004 | 39 | 0.05 | 0.005 |
| | | Outer Lagoon | 88 | 0.11 | 0.005 | 40 | 0.09 | 0.005 |

Table S19. Statistical analysis of linear mixed effect models of coral fitness scores.

| Response Variable | Subset Analysis | Fixed Effect | SS | df | F | *p* |
|---|---|---|---|---|---|---|
| Fitness Score (G * S * R) | *M. capitata* | Origin<br>**Destination**<br>**Origin * Destination** | 0.06<br>**8.06**<br>**1.51** | 1, 18<br>**1, 18**<br>**1, 18** | 0.93<br>**133.73**<br>**25.04** | 0.35<br>**<0.01**<br>**<0.01** |
| Fitness Score (G * S) | *M. capitata* | Origin<br>**Destination**<br>Origin * Destination | 0.08<br>**2.56**<br>0.18 | 1, 18<br>**1, 18**<br>1, 18 | 0.87<br>**29.17**<br>2.06 | 0.36<br>**<0.01**<br>0.17 |
| | *P. compressa* | Origin<br>**Destination**<br>Origin * Destination | 0.02<br>**1.39**<br>0.02 | 1, 18<br>**1, 18**<br>1, 18 | 0.18<br>**16.22**<br>0.28 | 0.68<br>**<0.01**<br>0.61 |

SS = Sum of Squares; df = degrees of freedom (numerator, denominator); F = F statistic; bold indicates statistical significance (p<0.05).

Table S20. Summary statistics of coral fitness scores.

| | | | **Fitness Score** | | | | | |
|---|---|---|---|---|---|---|---|---|
| **Species** | **Origin** | **Destination** | G * S * R | | | G * S | | |
| | | | N | mean | sem | N | mean | sem |
| *Montipora capitata* | Inner Lagoon | Inner Lagoon | 10 | 0.89 | 0.08 | 10 | 1.77 | 0.17 |
| | | Outer Lagoon | 10 | 2.17 | 0.15 | 10 | 2.41 | 0.17 |
| | Outer Lagoon | Inner Lagoon | 10 | 1.40 | 0.09 | 10 | 1.75 | 0.11 |
| | | Outer Lagoon | 10 | 1.91 | 0.09 | 10 | 2.12 | 0.10 |
| *Porites Compressa* | Inner Lagoon | Inner Lagoon | NA | NA | NA | 10 | 1.48 | 0.08 |
| | | Outer Lagoon | NA | NA | NA | 10 | 1.81 | 0.14 |
| | Outer Lagoon | Inner Lagoon | NA | NA | NA | 10 | 1.38 | 0.11 |
| | | Outer Lagoon | NA | NA | NA | 10 | 1.80 | 0.13 |

*N indicates parental colony*

Table S21. Linear mixed effect model statistical analyses of coral response variables following the acute heat stress experiment. Subset analysis indicates factors for which separate analyses were conducted.

| **Response Variable** | **Subset Analysis** | **Model Class** | **Fixed Effects** | **Random Effects** |
|---|---|---|---|---|
| Gross Photosynthesis | Time point | lmer [1] | Origin Site; Destination Site; Species | Parent |
| Net Photosynthesis | Time point | lmer [1] | Origin Site; Destination Site; Species | Parent |
| Respiration | Time point | lmer [1] | Origin Site; Destination Site; Species | Parent |
| Calcification | Time point | lmer [1] | Origin Site; Destination Site; Species | Parent |
| Dark Adapted Yield (log transformed) | Time point | lmer [1] | Origin Site; Destination Site; Species | Parent |

1. *lme4* package (Bates et al. 2015); Responses analyzed by Type III Satterthwaite ANOVA tests.

Table S22. Statistical analysis of linear mixed effect models of dark adapted yield following the acute heat stress experiment.

| **Response Variable** | **Subset Analysis** | **Fixed Effect** | **SS** | **df** | **F** | ***p*** |
|---|---|---|---|---|---|---|
| Dark Adapted Yield | Stress | Origin | 1.73e-04 | 1, 34 | 0.03 | 0.87 |
| | | Destination | 5.19e-05 | 1, 120 | 0.01 | 0.93 |
| | | **Species** | **8.83e-02** | **1, 34** | **14.34** | **<0.01** |
| | | **Treatment** | **8.19e-01** | **1, 130** | **132.98** | **<0.01** |
| | | Origin x Destination | 1.55e-03 | 1, 120 | 0.25 | 0.62 |
| | | Origin x Species | 7.24e-03 | 1, 34 | 1.18 | 0.29 |
| | | Destination x Species | 7.76e-03 | 1, 120 | 1.26 | 0.26 |
| | | Origin x Treatment | 3.98e-04 | 1, 130 | 0.06 | 0.80 |
| | | Destination x Treatment | 9.03e-05 | 1, 130 | 0.01 | 0.90 |
| | | **Species x Treatment** | **7.13e-02** | **1, 130** | **11.58** | **<001** |
| | | Origin x Destination x Species | 1.73e-04 | 1, 120 | 0.03 | 0.87 |
| | | Origin x Destination x Treatment | 1.26e-03 | 1, 130 | 0.21 | 0.65 |
| | | Origin x Species x Treatment | 2.19e-05 | 1, 130 | 0.00 | 0.95 |
| | | Destination x Species x Treatment | 3.06e-04 | 1, 130 | 0.05 | 0.82 |
| | | Origin x Destination x Species x Treatment | 9.39e-04 | 1, 130 | 0.15 | 0.70 |

SS = Sum of Squares; df = degrees of freedom (numerator, denominator); F = F statistic; bold indicates statistical significance ($p<0.05$). Data were log transformed to meet assumptions of normality. Homogeneity of variance was violated in analysis of dark-adapted yield.

Table S23. Statistical analysis of linear mixed effect models of coral metabolic rates following the acute heat stress experiment.

| Response Variable | Subset Analysis | Fixed Effect | SS | df | F | *p* |
|---|---|---|---|---|---|---|
| Gross Photosynthesis | Stress | Origin | 1.22e-04 | 1, 35 | 1.08 | 0.31 |
| | | Destination | 1.52e-04 | 1, 112 | 1.35 | 0.25 |
| | | **Species** | **5.69e-04** | **1, 35** | **5.04** | **0.03** |
| | | **Treatment** | **7.97e-03** | **1, 117** | **70.62** | **<0.01** |
| | | Origin x Destination | 2.39e-04 | 1, 112 | 2.12 | 0.15 |
| | | Origin x Species | 7.39e-05 | 1, 35 | 0.66 | 0.42 |
| | | Destination x Species | 3.29e-04 | 1, 112 | 2.92 | 0.09 |
| | | Origin x Treatment | 6.85e-05 | 1, 117 | 0.61 | 0.44 |
| | | Destination x Treatment | 4.76e-05 | 1, 117 | 0.42 | 0.52 |
| | | **Species x Treatment** | **7.67e-04** | **1, 117** | **6.80** | **0.01** |
| | | Origin x Destination x Species | 2.46e-04 | 1, 112 | 2.18 | 0.14 |
| | | Origin x Destination x Treatment | 1.35e-04 | 1, 117 | 1.20 | 0.28 |
| | | Origin x Species x Treatment | 0.00 | 1, 117 | 0.00 | 0.99 |
| | | Destination x Species x Treatment | 5.72e-05 | 1, 117 | 0.51 | 0.48 |
| | | Origin x Destination x Species x Treatment | 2.14e-04 | 1, 117 | 1.90 | 0.17 |
| Respiration | Stress | Origin | 3.24e-06 | 1, 37 | 0.17 | 0.68 |
| | | Destination | 2.32e-06 | 1, 112 | 0.12 | 0.73 |
| | | Species | 6.33e-05 | 1, 37 | 3.28 | 0.08 |
| | | **Treatment** | **3.80e-04** | **1, 117** | **19.67** 0.54 | **<0.01** |
| | | Origin x Destination | 1.05e-05 | 1, 112 | 0.23 | 0.46 |
| | | Origin x Species | 4.52e-06 | 1, 37 | 1.82 | 0.63 |
| | | Destination x Species | 3.52e-05 | 1, 112 | 0.15 | 0.18 |
| | | Origin x Treatment | 2.92e-06 | 1, 117 | 0.24 | 0.70 |
| | | Destination x Treatment | 4.62e-06 | 1, 118 | **10.83** 2.39 | 0.63 |
| | | **Species x Treatment** | **2.09e-04** | **1, 117** | 1.83 | **<0.01** |
| | | Origin x Destination x Species | 4.62e-05 | 1, 112 | 0.13 | 0.12 |
| | | Origin x Destination x Treatment | 3.54e-05 | 1, 118 | 1.02 | 0.18 |
| | | Origin x Species x Treatment | 2.52e-06 | 1, 117 | 2.05 | 0.72 |
| | | Destination x Species x Treatment | 1.97e-05 | 1, 118 | | 0.31 |
| | | Origin x Destination x Species x Treatment | 3.97e-05 | 1, 118 | | 0.15 |
| P:R Ratio | Stress | Origin | 1.73e-01 **1.25** | 1, 35 | 0.80 | 0.38 |
| | | **Destination** | 1.30e-04 **15.12** | **1, 111** | **5.79** | **0.02** |
| | | Species | 3.75e-01 | 1, 35 | 0.00 | 0.98 |
| | | **Treatment** | 2.06e-01 | **1, 116** | **69.85** | **<0.01** |
| | | Origin x Destination | 6.30e-03 | 1, 111 | 1.73 | 0.19 |
| | | Origin x Species | 1.96e-01 **1.71** | 1, 35 | 0.95 | 0.33 |
| | | Destination x Species | 2.85e-01 | 1, 111 | 0.03 | 0.86 |
| | | Origin x Treatment | 8.86e-03 | 1, 116 | 0.90 | 0.34 |
| | | **Destination x Treatment** | 1.13e-01 | **1, 116** | **7.91** | **<0.01** |
| | | Species x Treatment | 1.37e-03 | 1, 116 | 1.32 | 0.25 |
| | | Origin x Destination x Species | 1.61e-02 | 1, 111 | 0.04 | 0.84 |
| | | Origin x Destination x Treatment | 9.13e-02 | 1, 116 | 0.52 | 0.47 |
| | | Origin x Species x Treatment | | 1, 116 | 0.01 | 0.94 |
| | | Destination x Species x Treatment | | 1, 116 | 0.07 | 0.79 |
| | | Origin x Destination x Species x Treatment | | 1, 116 | 0.42 | 0.52 |

SS = Sum of Squares; df = degrees of freedom (numerator, denominator); F = F statistic; bold indicates statistical significance ($p<0.05$).

Table S24. Statistical analysis of linear mixed effect models of calcification following the acute heat stress experiment.

| Response Variable | Subset Analysis | Fixed Effect | SS | df | F | *p* |
|---|---|---|---|---|---|---|
| Calcification | Stress | Origin | 3.18e-01 | 1, 39 | 0.72 | 0.40 |
| | | Destination | 5.50e-02 | 1, 115 | 0.13 | 0.72 |
| | | **Species** | **17.68** | **1, 39** | **40.23** | **<0.01** |
| | | **Treatment** | **12.05** | **1, 119** | **27.41** | **<0.01** |
| | | Origin x Destination | 2.25e-01 | 1, 115 | 0.51 | 0.48 |
| | | Origin x Species | 4.22e-01 | 1, 39 | 0.96 | 033 |
| | | Destination x Species | 1.89e-01 | 1, 115 | 0.43 | 0.51 |
| | | Origin x Treatment | 3.64e-01 | 1, 119 | 0.83 | 0.36 |
| | | Destination x Treatment | 1.18 | 1, 120 | 2.68 | 0.10 |
| | | Species x Treatment | 1.64 | 1, 119 | 3.72 | 0.06 |
| | | Origin x Destination x Species | 1.08e-01 | 1, 115 | 0.25 | 0.62 |
| | | Origin x Destination x Treatment | 8.21e-01 | 1, 120 | 1.87 | 0.17 |
| | | Origin x Species x Treatment | 2.82e-01 | 1, 119 | 0.64 | 0.42 |
| | | **Destination x Species x Treatment** | **2.02** | **1, 120** | **4.59** | **0.03** |
| | | Origin x Destination x Species x Treatment | 2.61e-01 | 1, 120 | 0.59 | 0.44 |

SS = Sum of Squares; df = degrees of freedom (numerator, denominator); F = F statistic; bold indicates statistical significance ($p<0.05$).

Table S25. Microsatellite results for *Montipora capitata*. Each letter indicates a unique allele at that locus. Blank cells indicate no data, as not all loci amplified for all colonies.

| | **Primer loci** | | | | | | | |
|---|---|---|---|---|---|---|---|---|
| **Parent colony** | Mc0004 | Mc0067 | Mc0163 | Mc0701 | Mc0797 | Mc0872 | Mc0903 | Mc0947 |
| 21 | | | A | | | | | |
| 22 | A | A | B | A | | A | A | A |
| 24 | B | A | C | B | A | B | B | B |
| 51 | C | B | | B | B | B | C | |
| 52 | D | | | B | C | C | C | B |
| 55 | | | D | A | C | C | B | |
| 71 | | | | B | | | | |
| 72 | | | | A | | | | |
| 121 | | C | B | B | C | | A | |
| 125 | | D | | A | C | B | | |
| 143 | | E | B | B | D | B | B | |
| 144 | | D | B | C | | | | |
| 162 | | F | | | | | B | |
| 163 | | D | B | D | C | | A | B |
| 166 | | | B | B | A | B | | |
| 167 | | D | | | C | | A | |
| 169 | | E | | A | | | D | |
| 170 | | | | | | | D | |
